# Remembering the “when”: Hebbian memory models for the time of past events

**DOI:** 10.1101/2022.11.28.518209

**Authors:** Johanni Brea, Alireza Modirshanechi, Georgios Iatropoulos, Wulfram Gerstner

## Abstract

Humans and animals can remember how long ago specific events happened. Little is known about the neural mechanisms that enable remembering the “when” of memories stored for long durations in the episodic memory system – in contrast to interval-timing on the order of seconds and minutes. Based on a systematic exploration of neural coding, association and retrieval schemes, we develop model classes that span the space of possible mechanisms for the reconstruction of the time of past events. In concrete examples we show how network architecture, Hebbian plasticity, synaptic pruning or systems consolidation allow the retrieval of the time of past events. In a simulation, we demonstrate how these mechanisms would enable food caching animals like corvids to remember what they cached, where, and how long ago. To dissociate different hypotheses, we propose three kinds of novel, non-verbal experiments, that can be run with humans and animals. Our simulations predict the experimental results for different classes of models. Our study shows that remembering the “when” can be implemented by many biologically plausible mechanisms and that carefully designed experiments are needed to pin down the actual neural implementation of the memory for the time of past events in different species.

**Author summary:** Was it yesterday, a week ago, or perhaps even longer since I last ate pizza? Questions like these are typically easy to answer, because humans have a good sense of how long ago specific events happened. Research involving animals, such as food-caching birds, has revealed that they, too, can remember the “when” of past events. So far, little is known about the neuronal mechanisms that enable remembering the “when”. In this study, we explore systematically concrete hypotheses of how neural circuits, synaptic plasticity and ongoing restructuring of neural circuits allow remembering the “when”. We propose experiments, that could be done with humans or animals, to discriminate different biologically plausible mechanisms for remembering the “when”.

## 1 Introduction

Humans and animals track temporal information on multiple timescales, to estimate, for example, the location of a sound source based on millisecond time differences of sound arrival at the two ears, the interval duration between the perception of lightning and thunder, or the days, months and years that have elapsed since an autobiographical event took place [1–9]. For autobiographical memories, recall of the “when” information is often an explicit and conscious reconstruction-based process [10], for example, “we went to Turkey the year my sister got married, she is five years older than me, got married at the age of 30, and I am now 37 years old, so this must have been 12 years ago.” However, even without explicit reconstruction, healthy human adults usually have a good sense of whether a recalled event happened yesterday, a year ago or decades ago, and there is evidence for automatic processes, in particular in young infants without a fully developed episodic memory system [11–13]. Also corvids, rodents, and other species with an episodic-like “what-where-when” memory can remember the time of past events on timescales of days to months [13, 14].

Whereas on timescales from milliseconds to minutes, multiple mechanisms based on changing neuronal activity patterns are known to support accurate interval timing [4–9], little is known about the neuronal mechanisms that support the recall of the time of past events on much longer timescales. Memories on these timescales are likely to rely on long-term synaptic plasticity [15] and possibly on systems consolidation [16].

Multiple research communities have developed models of episodic memory to explain recall of past events [17]. These models focus on different aspects, like replicating data from laboratory-based memory tasks, such as free or serial recall of lists (reviewed in [18]), embedding episodic memory in cognitive architectures (reviewed in [19]), developing attractor neural networks consistent with anatomical and physiological knowledge of the hippocampal formation (reviewed in [20]), or explaining systems consolidation (reviewed in [16]). However, little work has been devoted to understand specifically how humans and animals remember the time of past events, although attempts at categorizing different theories of the time of past events have been made [10].

Here, we study from a theoretical perspective different neural mechanisms that enable *automatic* reconstruction of the time of past events on long timescales. Whereas our theoretical considerations about the representation of temporal information are relevant for tracking time on any scale, we focus, in particular, on settings used to investigate episodic-like memory, where a stream of sensory inputs on a timescale of days or months is perceived by an organism that can recall past events and respond with some actions (Figure 1A). As an example of the abstract setting described in Figure 1A, one may think of experiments where food caching animals learn to retrieve from caches they made the same day and ignore caches they made a few days ago (see Figure 1B and Table 1). For memory tasks on this timescale, it is unlikely that some kind of persistent neural activity bridges the gap between storage and recall. Instead, it is commonly believed that memories are stored over long timescales in an activity-silent manner: synaptic connections are altered through Hebbian plasticity mechanisms [15, 21] at the moment when an event is experienced. These connections are later reactivated by stimulating neurons upstream of the altered connections, thereby transforming dormant memories back into neural activity patterns. Our goal is to explore possible Hebbian mechanisms that allow to decode the age of such dormant memories at the moment of retrieval (Figure 1C) As the neural mechanisms underlying the retrieval of the age of dormant memories are largely unknown and may vary across different species, we do not want to limit ourselves, a priori, to specific classes of models; rather we aim at spanning the space of possible mechanisms. We discuss in detail some representative examples and propose in simulations specific behavioral experiments that allow to distinguish between different model classes. Importantly, and going beyond behavioral predictions, we also show that detailed neural and synaptic recordings are needed to identify the exact mechanisms that allow retrieval of the “when” of past events.

**Table 1.**
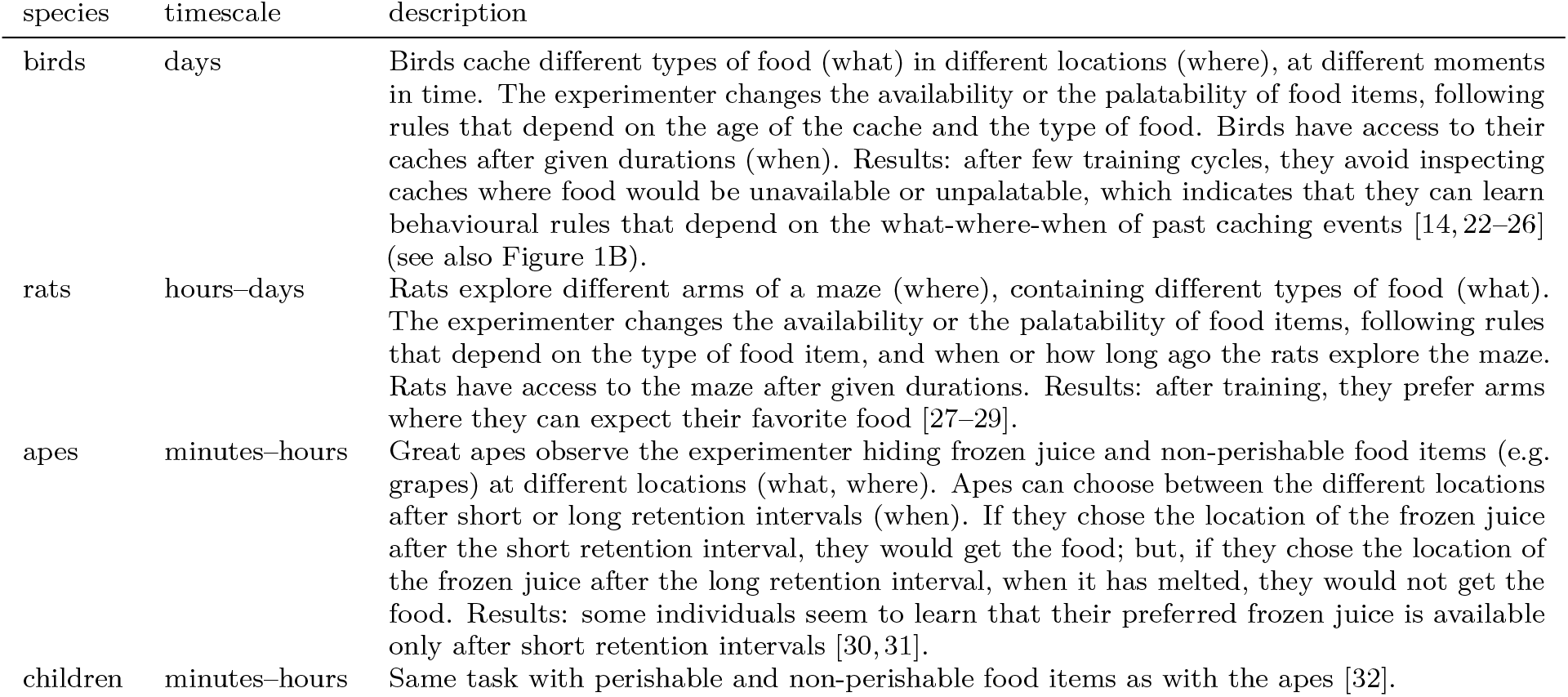
Examples of non-verbal behavioral experiments to investigate memory for the “when” of past events.

| species | timescale | description |
| --- | --- | --- |
| birds | days | Birds cache different types of food (what) in different locations (where), at different moments in time. The experimenter changes the availability or the palatability of food items, following rules that depend on the age of the cache and the type of food. Birds have access to their caches after given durations (when). Results: after few training cycles, they avoid inspecting caches where food would be unavailable or unpalatable, which indicates that they can learn behavioural rules that depend on the what-where-when of past caching events [14, 22–26] (see also Figure 1B). |
| rats | hours–days | Rats explore different arms of a maze (where), containing different types of food (what). The experimenter changes the availability or the palatability of food items, following rules that depend on the type of food item, and when or how long ago the rats explore the maze. Rats have access to the maze after given durations. Results: after training, they prefer arms where they can expect their favorite food [27–29]. |
| apes | minutes–hours | Great apes observe the experimenter hiding frozen juice and non-perishable food items (e.g. grapes) at different locations (what, where). Apes can choose between the different locations after short or long retention intervals (when). If they chose the location of the frozen juice after the short retention interval, they would get the food; but, if they chose the location of the frozen juice after the long retention interval, when it has melted, they would not get the food. Results: some individuals seem to learn that their preferred frozen juice is available only after short retention intervals [30, 31]. |
| children | minutes–hours | Same task with perishable and non-perishable food items as with the apes [32]. |

**Fig 1.**
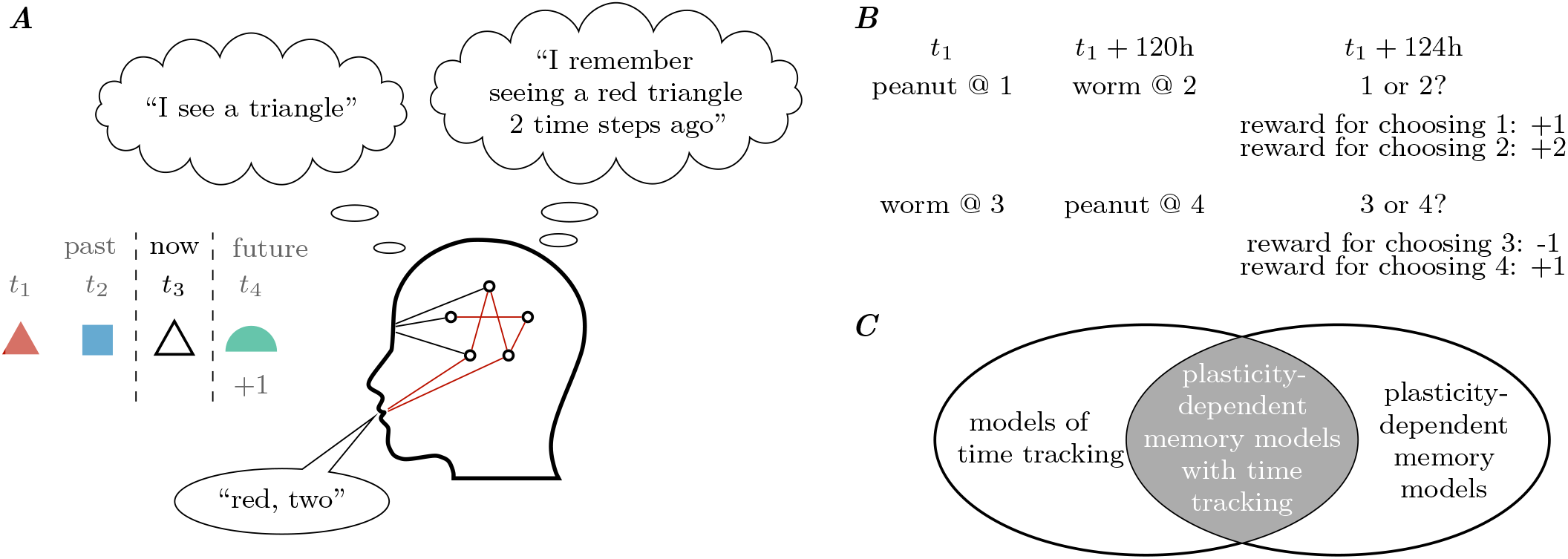
A simple setting to study episodic-like memory. **A** We consider discrete sensory streams, like perceiving a red triangle at time *t*_1_ and a blue square at time *t*_2_. In all illustrations we use color, shape and time as abstract analogies of the “what”, “where” and “when” of specific events, respectively. The time points *t*_1_, *t*_2_, … are not necessarily equally spaced and may be separated by hours or days. At time *t*_3_, the white triangle first triggers activity in the brain that corresponds to the perception of a white triangle and, second, activity that corresponds to remembering the red triangle, including the information of how long ago the red triangle was perceived. An action is performed upon memory retrieval, like saying “red, two”. In the next time step, a new stimulus can be given, together with a reinforcement signal (+1). Our goal is to find neural network dynamics and synaptic plasticity rules that change the connections between neurons (red lines) such that the sensory stream can be remembered, and action selection rules that depend on the recalled event can be learned. **B** For example, food caching birds can learn to retrieve caches with fresh worms, and ignore caches with old worms [33]. In trials, where they cache peanuts first (at a given location, indicated by “@ 1”), and worms 120h later, they can retrieve fresh worms (which they prefer over peanuts; indicated with reward +2). In trials where they cache worms first, the worms are unpalatable at retrieval (indicated by reward −1), but the peanuts are still fresh. Clayton et al. [33] showed convincingly that the birds’ ability to learn to ignore caches with old worms depends on memory of the “what” and “how long ago” of caching events. **C** Many models of time tracking in the milliseconds to seconds range, such as pacemaker-accumulator models or state-dependent networks [6], do not depend on (long-term) synaptic plasticity, but on changing neural activity. Most existing models of plasticity-dependent memory, such as models of the hippocampus [20], focus on storage and retrieval of memories (the “what” and the “where”), but not on the age of memories (the “when”). In this work, we investigate plasticity-dependent memory models that keep track of time on long timescales from days to years (indicated by the gray area).

## 2 Results

### 2.1 Representing Information: the Space of Possible Codes

The activity in a network of neurons can represent a memory in multiple ways. For simplicity, we assume that the “what”, the “where” and the “when” of each memory are elements of discrete sets, like the sets of colors, shapes and time points in Figure 1A. The value of such discrete variables can be represented with, (i) the firing rate of a neuron (`rate`), (ii) the identity of an active neuron within a group of neurons (`onehot`) or (iii) the distributed activity pattern in a group of neurons (`distr`; see Figure 2A and section “Formal Description of Codes”). For example, in a rate code of color, the sight of red and blue objects evokes different activity levels in the same neuron, whereas in a one-hot code, different neurons are tuned to different colors. A strict rate code with a single neuron or a strict one-hot code, where a given stimulus feature activates a single neuron, are idealizations that are unlikely to be found in any brain. Instead, stimulus features may be represented by a distributed code, where multiple neurons become active, when perceiving the color “red”, for example. However, certain distributed codes can be reduced to rate or one-hot codes by summing the activity of subsets of neurons. Trivial examples are redundant rate or one-hot codes with groups of identical neurons. Another example is the population rate code (`poprate`), where the number of active neurons encodes the value of a variable. We use the term “distributed code” only when such a reduction by summation is impossible.

**Fig 2.**
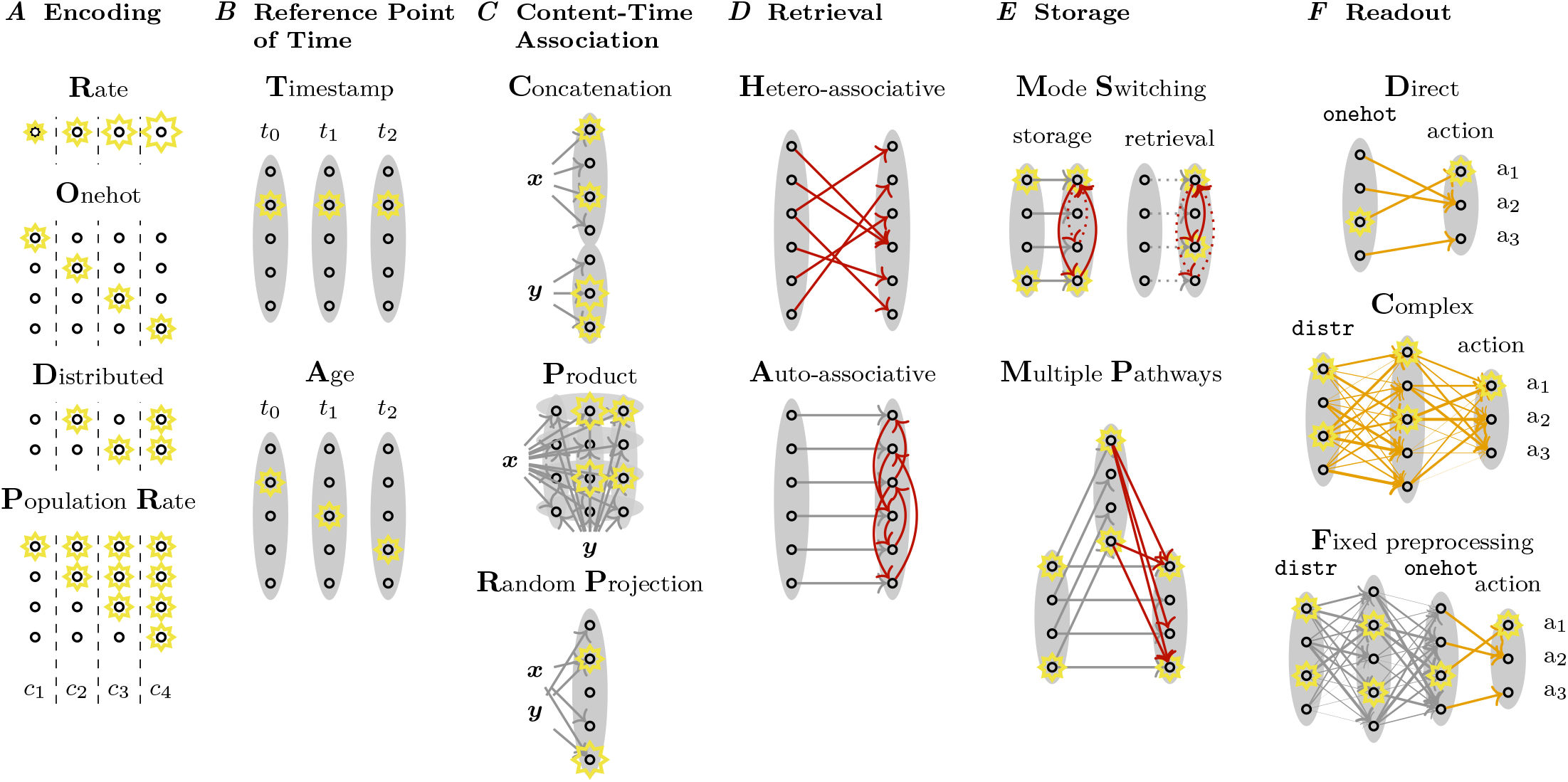
Several distinct “what-where-when” memory systems can be constructed by combining specific choices of the encoding, reference point of time, association, retrieval, storage and readout schemes. **A** The elements *c*_1_, *c*_2_, *c*_3_, *c*_4_ of a set *C* can be encoded, for example, (from top to bottom) in a rate code, one-hot code, distributed code or a population rate code (see Formal Description of Codes). **B** In timestamp memory systems, time is measured relative to a fixed reference point and the “what-where-when” information of a specific event is, therefore, encoded with the same activity pattern at storage *t*_0_, and at any time of recall *t*_1_ or *t*_2_. In age memory systems, the retrieved memory pattern of a given event changes with time, because time is measured relative to a changing reference point. **C** Information ***x*** and ***y***, about e.g. the content and the time of an event, can be associated in multiple ways; for example, with concatenation, or non-linear mixed codes, like the (outer) product, or random projection code. The two-dimensional arrangement of the twelve neurons in the product code is for visualization purposes; they could also be arranged in a vector with 12 elements. **D** Memory retrieval can be based on hetero-associative recall with learned feed-forward synaptic connections (red) or on auto-associative recall with learned recurrent synaptic connections (red). **E** Separation of storage and retrieval can be achieved with external modulators that switch the network mode from being input-driven in storage mode, to being recurrently driven in retrieval mode. Alternatively, multiple pathways allow a separation of storage and retrieval through different signal propagation properties along the different pathways. **F** The difficulty of learning flexible rules depends on the code. Direct readout: For one-hot coding, any rule is learnable with direct connections to action neurons. Complex readout: Learning arbitrary rules based on distributed or rate coding can be achieved with plastic connections in multilayer perceptrons or complex readout with fixed preprocessing: with hard-wired transformations into, for example, one-hot codes that allow flexible learning. See Table 2 for an overview of different models that can be constructed with different combinations.

#### 2.1.1 Representing Time: Timestamp and Age Codes, Internal and External Zeitgeber

Information about the “when” of an event can be represented by any code discussed above, as soon as a reference point for measuring time is defined. We distinguish timestamp and age representations (Figure 2B). In *timestamp* representations, time is measured relative to a fixed reference point in time. The fixed reference point could be the birth of an individual and the “when” information of an event could be represented as “5 months since birth”. In timestamp representations of time, the neural activity representing the “when” information during recall of a given event is always the same, no matter when recall happens; this neural activity code can thus be seen as representing a timestamp attached to each memory. Importantly, we do not assume that this timestamp representation encodes literally the date and time of an event. In fact, any neural activity pattern can be a timestamp, if it allows to infer the time of a given event and does not change with the age of the memory. In contrast, in *age* representations, time is measured relative to changing moments in time. The changing reference point could be the current moment in time and the “when” information of an event could be represented as “8 months ago”. In contrast to timestamp representations of time, age coding implies that the neural activity during recall of a given event is not the same at different moments, because the “when” information depends on how much time has elapsed between storage and recall; this neural activity code can thus be seen as representing the age of each memory. The distinction between timestamp and age coding is similar to an important difference in McTaggart’s A and B series [34]: in age coding the representation of the “when” of an event changes continually, like the position of an event in the A series, whereas in timestamp coding the representation is constant, like the position of an event in the B series. An example of an age code is shown in Figure 2B, where the elapsed time between storage and recall is represented by the location of the activity peak.

**Table 2.**
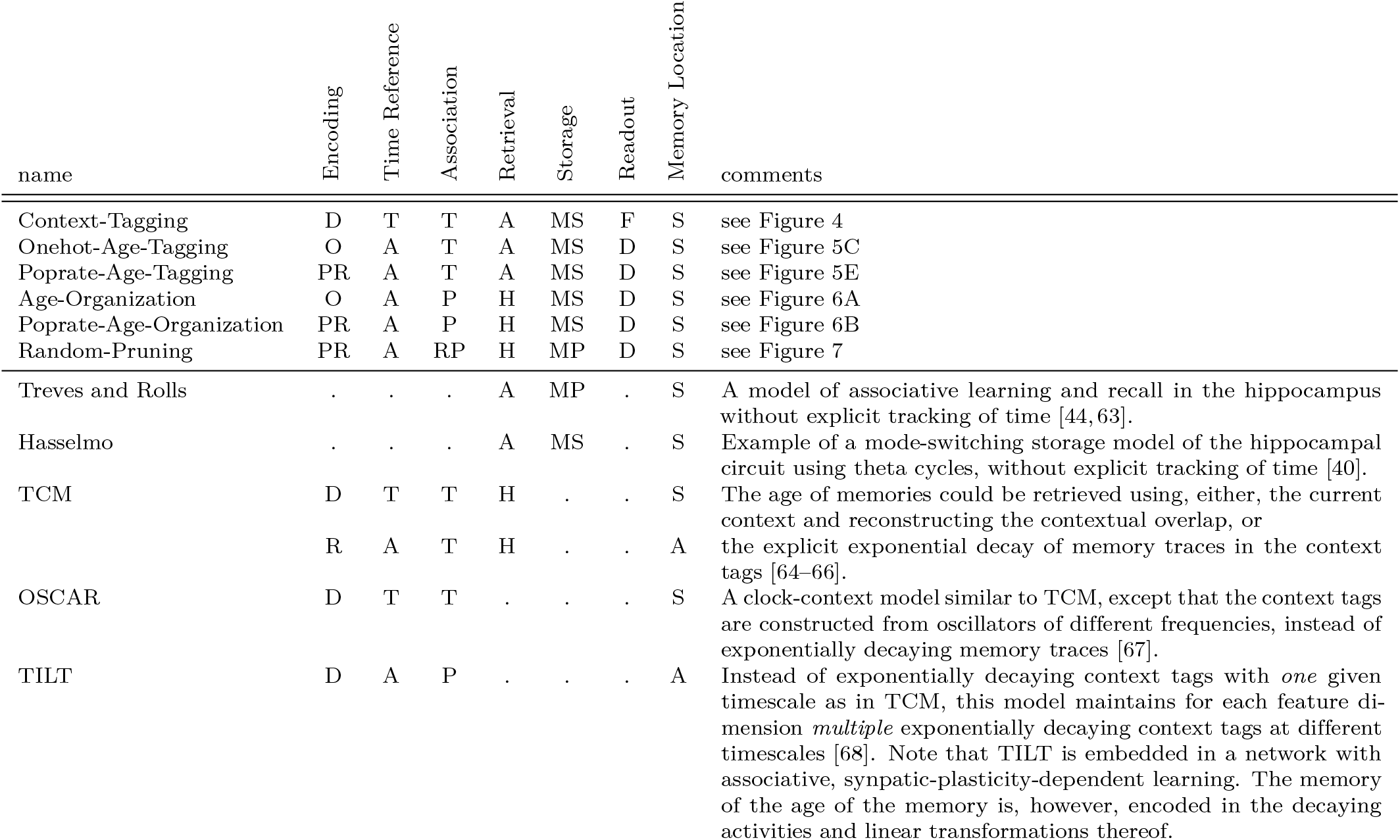
Comparison of Computational Models. The labels in columns “Encoding” to “Readout” correspond to the labels used in Figure 2. The label used in column “Memory Location” indicates whether the memory is stored synaptically (S), or in the activity pattern (A) of some neurons. A dot “.” indicates that the model is agnostic about this feature.

We use the term *zeitgeber* to refer to the process that generates either timestamps or changes the age code. Unlike a clock, that is synchronized with physical time, a zeitgeber may drive the representation of time with variable speed that depends, for example, on the frequency of events that are worth to be memorized. The zeitgeber can either be an internal process that runs almost autonomously inside the time-perceiving agent or it can depend mostly on the agent’s interaction with the external world. Internal zeitgebers can be any biological process inside the agent with a fairly stable time constant such as ramping or decreasing synaptic strengths, neurogenesis, spine turnover, the circadian rhythm (in the absence of exposure to sunlight) or even changes in satiety, thirst or tiredness level. Examples of external zeitgebers are processes like the ticking of a clock, the day-night cycle or the change of seasons; also processes that involve the agent’s actions, like changes of context (leaving or entering a house) or changes of the main activity (switching from working to eating lunch), could act as external zeitgebers.

### 2.2 Associating Information: How to Combine What, Where and When

The ability to remember everything about a given event, the “what”, “where” and “when” information needs to be associated in some way. In the following, we assume the “what” and the “where” are given as a content variable in some code and focus exclusively on how the content is associated with the “when” information. The question of associating the “what” (and “where”) with the “when” becomes the question of building an “association function” that produces a neural activity pattern in response to the “when” information on one side and the encoded content (“what” and “where”) information on the other side (see section “Formal Description of Association Schemes”).

The number of possibilities to write down such an association function is huge, even if we restrict ourselves to those functions that do not “lose” any information, in the sense that the content and “when” information can be faithfully reconstructed from the neural activity pattern during recall. In the following we focus on three specific examples of association functions: concatenation, (outer) product and random projection codes (Figure 2C).

In a *concatenation* code, neurons can be split into two separate groups: one representing the “when” information and the other one the content information (c.f. Figure 2C). Closely related is a *linear mixed code*, where such a split is not directly possible, because single neurons contribute to the representation of both content and “when” information, but a linear transformation of the neural population activity would allow to represent the content and “when” information in a concatenation code. An example of a *non-linear mixed code* is the *product code*, where the neural population activity is given by the outer product of the content and the “when” code. This product code is a special case of tensor product variable binding [35]. Such a product code requires, in general, more neurons than a concatenation code: if content and “when” information could be represented separately by *N* and *M* neurons, respectively, their association with a product code requires *N* × *M* neurons, whereas *N* + *M* neurons would be sufficient for a concatenation code. Other non-linear mixed codes can be constructed with (circular) convolutions [36, 37] or *random projections*, where the activity of each neuron in a group depends non-linearly on a randomly weighted mixture of content and “when” information.

The way a neuronal population represents the association of content and “when” information has important implications for the readout of retrieved memories, as we will discuss in the next section and demonstrate in the section “Experimental Predictions“.

### 2.3 Storage, Retrieval and Readout of Memories

So far, we considered only the representation of information. However, the description of a memory system is incomplete without a characterization of the storage, retrieval and readout mechanism.

#### 2.3.1 Retrieval

In the field of neural networks, memory retrieval is typically implemented with hetero- or auto-associative memories (Figure 2D). In both, hetero- and auto-associative networks, the output activity of a neural network in response to an input cue represents the retrieved memory. In a hetero-associative memory, retrieval is performed in a single step whereas a recurrent auto-associative network requires convergence to a fixed point [38]. However, a single update step is often sufficient to almost reach the fixed point and retrieve a memory almost perfectly, in particular in kernel memory networks [39].

#### 2.3.2 Separation of Storage and Retrieval

Suppose “red triangle” has already been stored at time step *t*_1_. At time *t*_3_, the input “white triangle” triggers recall of the stored memory, i.e. the neural code for the “when” information *t*_1_ together with the content information “triangle” and “red” should be accessible at time *t*_3_ and become active while the remembered event “red triangle” is retrieved from memory (retrieval phase). At the same time step *t*_3_, however, it must also be possible to store the new event “white triangle”, as an event that happens at time *t*_3_ (storage phase).

We distinguish two basic mechanisms to separate storage and retrieval: *mode switching* and *multiple pathways* (Figure 2E). A prominent example of mode switching is the Hasselmo et al. model [40], of a periodic process in which external input drives the memory network during the storage phase and recurrent connectivity in the memory network dominates during the retrieval phase. The theta rhythm in the hippocampus, or some modulatory factors, like neurotransmitters, could drive such a periodic process [40, 41], although the plausibility of this mechanism across species is debated [42, 43].

Alternatively, the separation of storage and retrieval could be implemented with separate pathways. For example, Treves and Rolls [44] suggested that the strong connections of the mossy fiber system from dentate gyrus (DG) to the CA3 hippocampal network clamp the CA3 neurons to an input-driven activity pattern during storage, whereas the perforant path to CA3 is used to initiate the retrieval process. In hetero-associative retrieval, strong clamping may not be needed, but different signal transmission delays along different pathways may be sufficient to separate storage from retrieval: if multiple pathways exist between two groups of neurons, for example the “input-content” pathway and the “input-intermediate-content” pathway in Figure 3, and if information travels at different speeds along the different pathways, then stimulus-driven activation through one pathway could be used for storage (“input-content” pathway in Figure 3) and through the other pathway for retrieval (“input-intermediate-content” pathway in Figure 3). In the simulated models in section “Experimental Predictions” we use this mechanism to distinguish storage and retrieval phases. In contrast to mode switching with 4-8 Hz theta frequency, multiple pathways allow for a quicker alternation between storage and retrieval on a timescale of few milliseconds. If plasticity were to be induced only during the storage phase, this would require mechanisms that are rather precisely timed. It remains to be seen, whether this is possible with fast modulation of Hebbian learning rules with a third factor [45–49]. However, ongoing plasticity may not be harmful, in particular with prediction-error reducing learning rules [50, 51].

**Fig 3.**
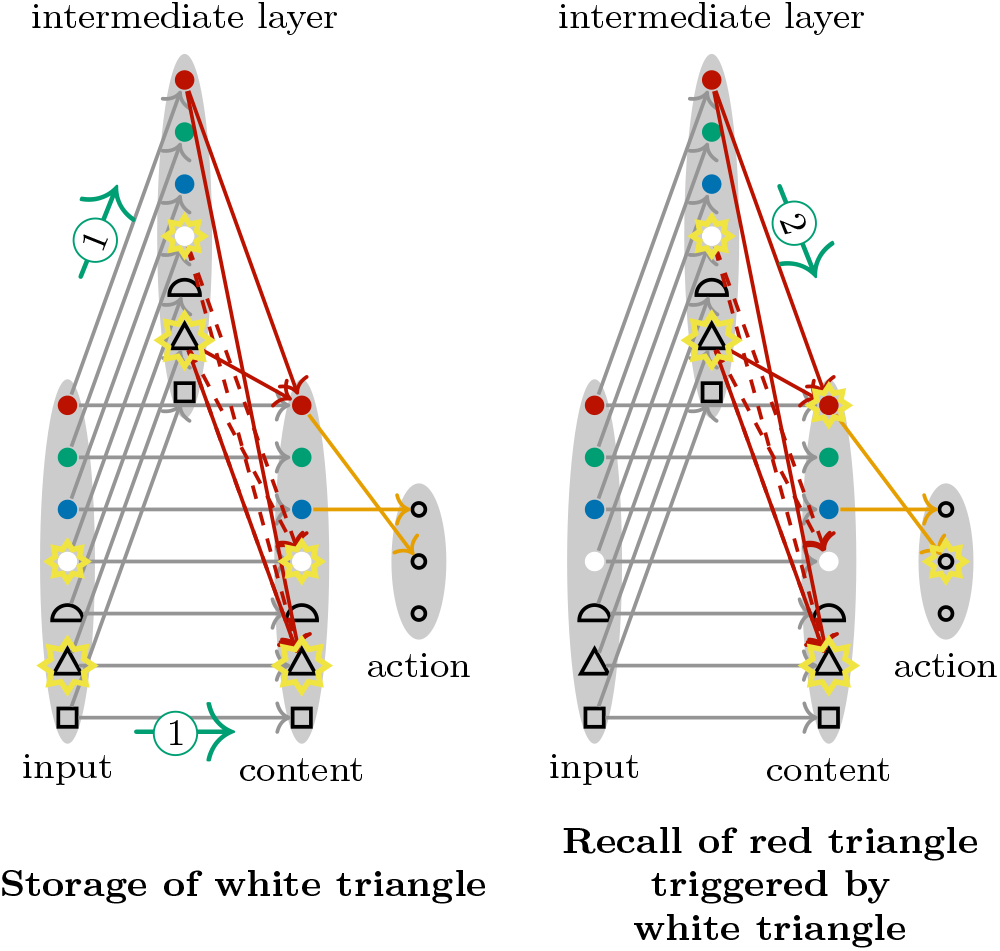
Example of a Storage and Retrieval Mechanism with Synaptic Delays. Signals take more time to travel in long pathways with multiple intermittent synapses than in short pathways, because signal transmission across chemical synapses takes time. Therefore, multiple pathways of different lengths between two groups of neurons can be used to separate storage and retrieval phases. During storage, the input drives the activity in the intermediate layer; the content layer receives input through the direct “input-content” pathway (indicated by green arrows with label 1). To simplify the exposition, the outgoing connections of the content layer are one-to-one in this example, but the same principle works in less artificial settings with random connectivity (see Figure 7). Connections between the intermediate and the content layer (red dashed arrows) are selected for growth with a Hebbian plasticity rule. Shortly thereafter, because of more synaptic delays along the longer pathway, the content neurons receive input through the “input-intermediate-content” pathway, i.e. the intermediate layer is the main input of the content layer (green arrow with label 2). Already grown connections (red arrows) enable recall of previous events. The actual growth of the connections selected in the storage phase (dashed red arrows) is not instantaneous and does therefore not interfere with the recall phase. Once the content neurons received input through the “input-intermediate-content” pathway, the content layer drives the action selection through weights that implement some learned rule (orange arrows).

#### 2.3.3 Behavioral Readout

Once a previously stored activity pattern is retrieved, it can trigger some behavioral output. In contrast to computer memory, where recall success can be measured by the number of bits lost between storage and recall, successful retrieval of a memory in humans and animals is usually inferred from some behavioral output, which may have a different representation than the input that led to the formation of the memory (e.g. visual input and vocal output in Figure 1A). Therefore, we must include a discussion of action selection that subjects perform in response to retrieved memories.

Behavioral rules control which action to perform in response to a specific retrieved memory. Whereas learning is often irrelevant for memory tasks with humans – as the behavioral rule is usually instructed – it is crucial for memory experiments with animals. How easily different behavioral rules can be learned depends on the representation of recalled memories. This can be used to design experiments that discriminate between different kinds of “what-where-when” memory systems, as we will show in section “Experimental Predictions”. For a strict one-hot code, for example the product of one-hot codes `onehot`(content) ⊗ `onehot`(age), any behavioral rule that maps content and age of a recalled event to a given action can be learned with *direct readout*, i.e. plastic connections between the layer of recalled activity to action neurons (Figure 2F). For distributed or rate coding, direct readout allows learning of some rules, but *complex readout* is needed to learn any rule. For example, the notorious XOR rule [53], where a certain action is taken if and only if two input neurons are jointly active or jointly inactive, cannot be learned with direct readout, but it can be learned with a multilayer perceptron (Figure 2F). Learning all connections in a multilayer perceptron can be achieved with the backpropagation algorithm or biologically plausible variants thereof [54–56], but it is rather slow if learning happens in an online fashion, where each example is used just once. An alternative is to rely on fixed weights in most layers to transform the input into a useful feature representation and quickly learn flexible mappings with biologically plausible Hebbian plasticity in the last layer (Figure 2F).

#### 2.3.4 Synaptic Plasticity

Long-term synaptic changes are presumably involved for memorization in the storage phase, for learning actions to indicate successful retrieval and to reflect the passage of time in age representations of time.

If pre- and postsynaptic neurons are jointly active during the storage phase, one-shot Hebbian synaptic plasticity (e.g. [57]) is sufficient to memorize and generate a trace of the event in the memory system. Behavioral rules that depend on the content and the age of recalled memories could be learned with neoHebbian synaptic plasticity [45–49], where jointly active neurons generate an eligibility trace that is modulated by a signal that arrives up to a few seconds later. This modulating signal could communicate the reward received after a successful action.

For age representations of time, synaptic turnover (growth or decay) could reflect the passage of time [58–62]. If the rate of synaptic pruning depends on the postsynaptic neuron, neurons with a low pruning rate would be active when young and old memories are retrieved, whereas neurons with a high pruning rate would only be active when young memories are retrieved. This would allow for a straightforward readout of the age of memories, as demonstrated in the examples below. A broad distribution of neuron-dependent pruning rates over timescales from hours and days to months and years would enable approximate recall of the age of memories (see also Figure 8). While such a neuron-dependent pruning rate is simple and plausible, we are not yet aware of clear experimental support for it.

For timestamp representations of time, the synaptic changes for memorization and behavioral learning are sufficient. Although this is an appealing advantage of timestamp representations of time, it comes at the cost of an increased complexity to compute the age of recalled memories, because a representation of the current moment in time needs to be compared with the time of storage of the recalled event.

### 2.4 Examples of Episodic-Like Memory Systems

With four different encoding schemes (Figure 2A), two different ways of representing time (Figure 2B), three different codes for associating content and time (Figure 2C), two different memory retrieval mechanisms (Figure 2D), three different readout architectures (Figure 2F), and two different storage-retrieval mechanisms (synaptic delays Figure 3 or periodic processes, like e.g. proposed by [40]) we have 4 × 2 × 3 × 2 × 3 × 2 = 288 concrete hypotheses about Hebbian “what-where-when” memory systems. This is a lower bound, because even more association, storage-retrieval and readout mechanisms are conceivable. The number 288 looks daunting. However, in this section we discuss in more details six specific examples that are representative of the different kinds of models (Table 2). As storage, retrieval and readout are general aspects of synaptic memory systems, irrespective of time, we selected our examples primarily to highlight advantages and disadvantages of different encodings of time, content-time associations, and timestamp versus age organization of time. Detailed mathematical descriptions of these models can be found in section “Mathematical Description of the Models“.

#### 2.4.1 Timestamp Tagging with Auto-Associative Retrieval and Complex Readout

Time tagging models are characterized by a concatenation code that combines one subgroup of neurons that represent content information (“content” in Figure 4) with another subgroup of neurons that represent “when” information (“tag” in Figure 4).

**Fig 4.**
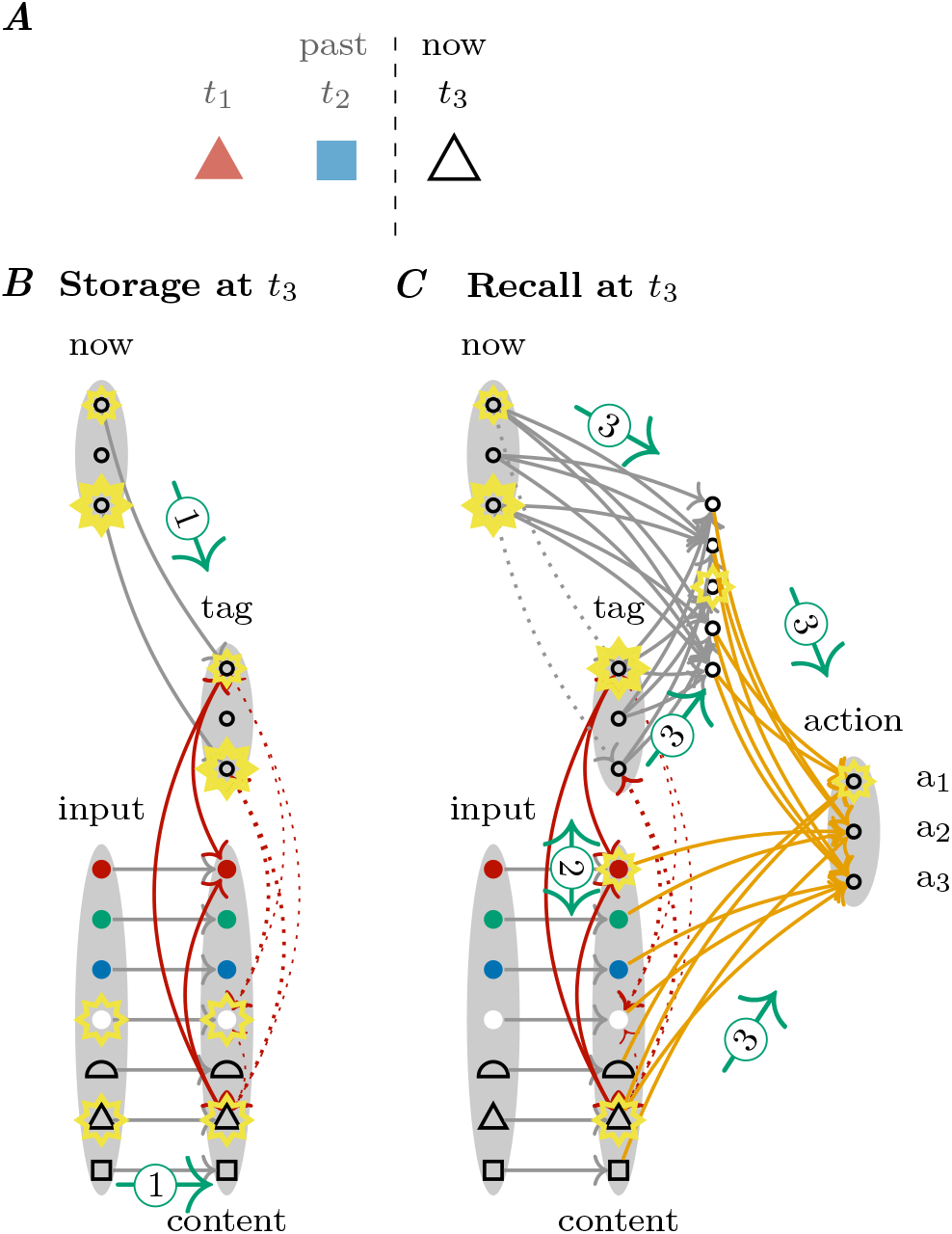
The Context-Tagging model: Timestamp Tagging with Auto-Associative Retrieval and Complex Readout. **A** The events “red triangle” and “blue square” were observed at times *t*_1_ and *t*_2_, respectively. At time *t*_3_ the event “white triangle” is observed. **B** During storage, the input drives the content layer, and the current context (activity in layer “now”) drives the “tag” neurons (green arrows with label 1) such that content and context can be bound together (dotted red lines). **C** Subsequently, the previously grown synaptic connections (red lines) allow auto-associative recall of the event “red triangle”. During recall, the tag neurons are no longer driven by input from the “now” neurons, but, through auto-associative recall (green arrow with label 2), the activity of the tag neurons encodes the context at time *t*_1_. The comparison of the current context (represented by the “now” neurons) with the recalled context (represented by the “tag” neurons) allows the readout network to estimate the age of the retrieved memory (green arrows with label 3). This comparison could be implemented in different ways, e.g. as a vector subtraction, or in a similar way as in linear prediction error networks [52]. Consequently, behavioral rules that depend on the age of the recalled memory can be learned (orange weights).

In the spirit of (temporal and random) context models [64–66], the “when” information could be given implicitly by activity patterns that encode context (“now” in Figure 4). This context could be a trace of recent observations of states that change on different timescales like emotional states, the presence of certain conspecifics, ambient temperature or the weather. On an implementation level and for timescales from seconds to hours, it could be given by so-called hippocampal time cells [9, 69–74] that fire at successive moments in temporally structured experiences. In the following, we call this the “Context-Tagging model”. At the moment of storage, this context information is bound together, e.g. by a Hebbian plasticity rule, with the specific event under consideration (Figure 4B). If an event triggers the recall of an earlier event (Figure 4C), the associated context, i.e. the “when” information, is also recalled. A comparison of the current context with the recalled context may allow a rough estimate of the age of the recalled memory (cf. contextual overlap theory, [10]). Because the estimation of age from the comparison of two context-related activity patterns does, in general, not induce a linearly separable problem, a complex readout network with at least one hidden layer is required, to learn arbitrary readout rules.

#### 2.4.2 Age Tagging Models

Other tagging models can be constructed with one-hot coding or population-rate coding for the age of memories (Figure 2A). In the Onehot-Age-Tagging model, storage leads to the formation of a synaptic connection to the first tag neuron (Figure 5A and Figure 5B). It is hypothesised that specific circuits allow to change the representation of the memory by growing new synapses and pruning old ones [75, 76]. Such a mechanism could implement a one-hot time code, where the identity of the activated tag neuron during recall indicates the age of the memory (Figure 5C).

**Fig 5.**
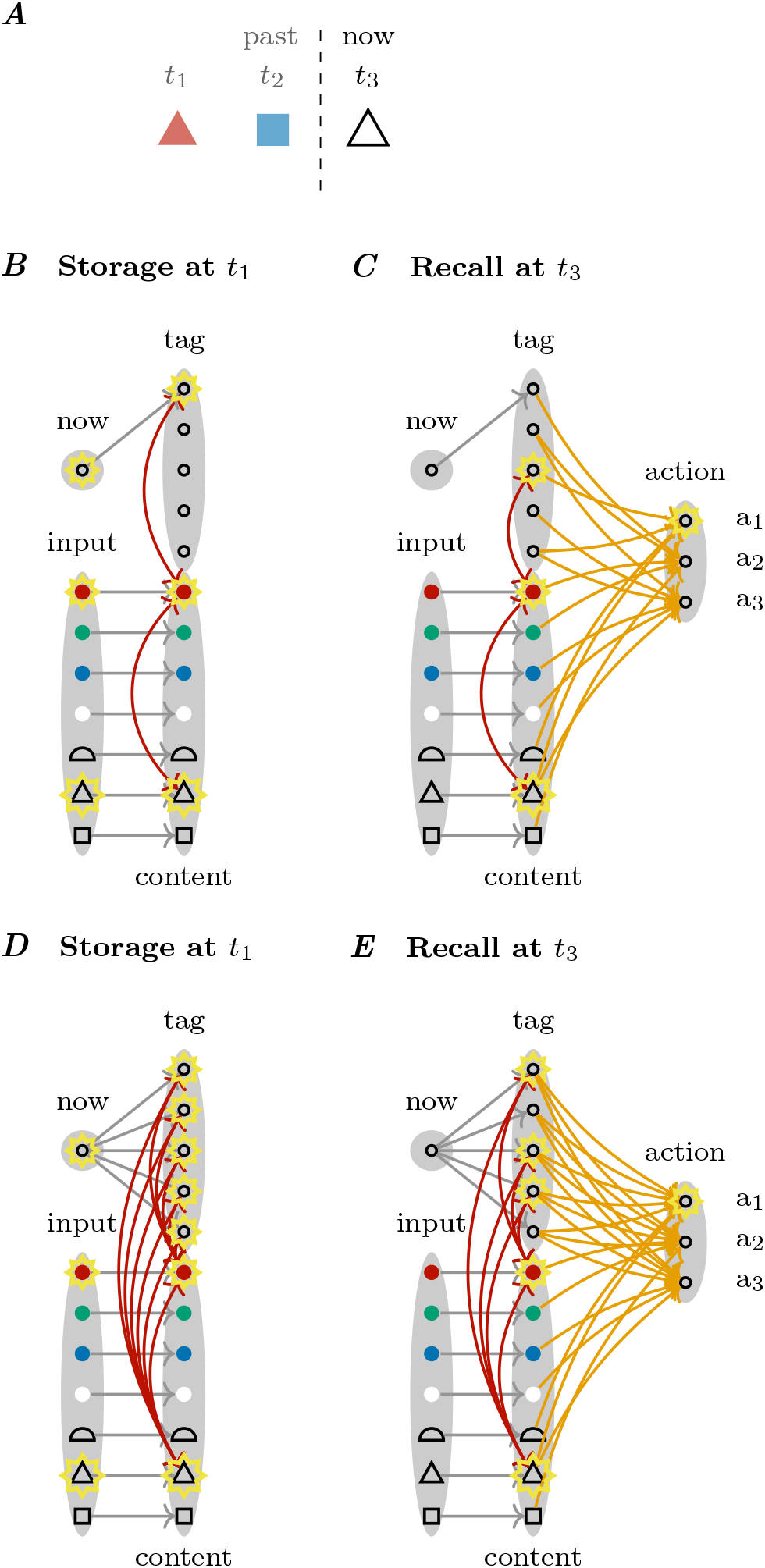
Age tagging models. **A** We consider the same sequence of events as in Figure 4A. **B** In the **Onehot-Age-Tagging** model, the first tag neuron is activated by the “now” neuron, which is only active during the storage phase, such that Hebbian one-shot learning establishes a connection between the first tag neuron and the active content neurons, when event “red triangle” is perceived at time *t*_1_. **C** Thanks to a process that prunes some synapses and grows new ones, tag neuron 3 is activated during recall at time *t*_3_, indicating how long ago the red triangle was observed. **D** In the **Poprate-Age-Tagging** model, connections to all tag neurons are formed during storage. **E** These connections are pruned at different moments in time, such that during recall at time *t*_3_, fewer connections are present than at time *t*_1_. The number of active “tag” neurons during recall encodes the elapsed time since storage: if many “tag” neurons are active during recall, the recalled event happened recently and if few “tag” neurons are active, it happened long ago.

In the Poprate-Age-Tagging model, storage leads to the formation of many synaptic connections to several “tag” neurons (Figure 5D). These connections are pruned at different moments in time. Therefore, the number of activated tag neurons during recall is indicative of the elapsed duration between storage and recall, and a simple readout mechanism that compares the sum of tag neuron activities to a fixed threshold – each action neuron in Figure 5E may have a different threshold – can implement behavioral rules that depend on the age of memories. We emphasize, however, that certain rules are impossible to learn with this simple readout mechanism (see e.g. Figure 11).

In contrast to the Context-Tagging model with a timestamp code (Figure 4), the representation of time changes in the Onehot-Age-Tagging model and the Poprate-Age-Tagging model, because the activity pattern of the tag neurons during retrieval depends on the elapsed time since storage. Age tagging allows simple readout learning of age-dependent behavioral rules, in particular, when the order of pruning synaptic connections is fixed, i.e. whenever it is possible to order pairs of tag neurons *i* and *j*, such that connections to tag neuron *i* are consistently lost earlier than simultaneously grown connections to tag neuron *j*. This could be implemented with neuron-dependent pruning rates as described in section Synaptic Plasticity. One can even prove (see Equivalence of One-Hot coding and Deterministic Population-Rate coding) that this population rate code leads to the same action selection policy as a model with one-hot coding of “when” information, if the readout connections follow a special synaptic plasticity rule.

#### 2.4.3 Age Organization with Systems Consolidation

Instead of using concatenation, as in tagging models, the content and “when” information could be associated with the (outer) product operation (c.f. Figure 2). Variants of models constructed in this way include the chronological organization of memories [10], which is also known as a shift register in engineering [77]. For example, this approach is implemented in the TILT model [68], an activity-based model of time-tracking memory (see also Table 2).

The chronological organization is most obvious, when arbitrarily encoded content information and one-hot encoded age information is associated with the product operation. We call this the Age-Organization model. In this case, the configuration of active neurons during recall of a given event depends on the time of recall (Figure 6), similarly to how suitcases on a conveyor belt change their position relative to a fixed observation point. Because of the product operation, there are multiple groups of neurons that code for content (e.g. the groups Δ*t*_0_, Δ*t*_1_, Δ*t*_2_ in Figure 6A), but the neurons in only one of these groups become active during recall of a specific event. The identity of the active group encodes implicitly the age of the memory: if recall happens some time interval Δ*t*_1_ after storage, the content neurons in group Δ*t*_1_ become active, whereas the neurons in other groups become active during recall at other times (Figure 6A).

**Fig 6.**
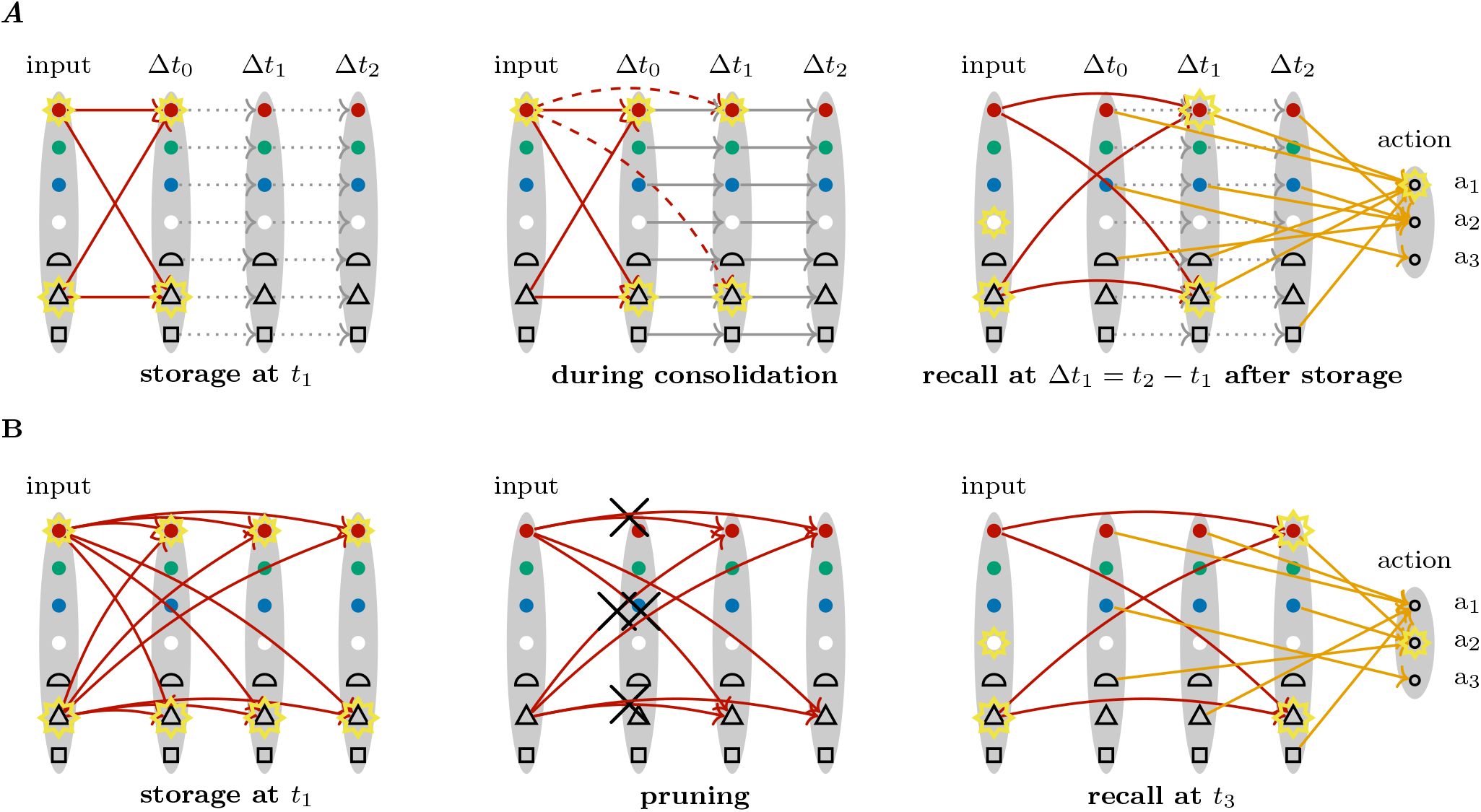
Examples of Chronological Organization Models. **A** In this age model, a systems consolidation mechanism shifts the location where a memory is stored. During storage (left), the red connections are strengthened, whereas the feedforward weights (gray dotted arrows) are inactive. During consolidation (middle), input neurons are randomly active and activity is forward propagated (gray arrows; indirect pathway from input to Δ*t*_1_), such that new, direct-pathway connections (dashed red) can grow between input and Δ*t*_1_ neurons. Simultaneously, the original weights between input and Δ*t*_0_ neurons decay, such that after consolidation only the newly grown weights remain. During recall (right), all active neurons in layers Δ*t*_0_, Δ*t*_1_, Δ*t*_2_, … give input to the action neurons (orange connections). **B** Another age model relies on pruning of synapses at different moments in time, similar to the Poprate-Age-Tagging model (Figure 5E). Synapses onto neurons in the first memory layer have a faster decay rate than those onto the last layer. During recall, the number of active neurons across all layers is indicative of the age of a memory: at *t*_1_ more “red” and “triangle” neurons are activated than at *t*_3_.

In contrast to activity-based models of time tracking such as the TILT model [68], a plasticity-dependent model of time tracking with an age code requires growing and pruning of synaptic connections. Similarly to the Onehot-Age-Tagging model, this could be mediated by a system’s consolidation process, where new synapses are grown to groups of neurons that code for older memories. For example, in consolidation phases during sleep, randomly activated input neurons could trigger recall of past events in neurons connected to the input by an indirect pathway, thereby allowing to create new direct-pathway connections (Figure 6A). A concrete example for how this could be implemented is given by the parallel pathway theory of systems consolidation [76]. The idea that the location of memorized events changes over time appears also in the multiple trace and trace transformation theory [16, 78]. The readout connections do not need to change during systems consolidation, but they allow to learn flexible rules, such as “perform action *a*_1_ if the memory for a red object is younger than Δ*t*_1_, and *a*_2_, if it is older than Δ*t*_2_, whereas for blue objects, perform action *a*_3_, if they are younger than Δ*t*_1_, and action *a*_2_, if they are older than Δ*t*_1_” (see orange lines in Figure 6A). If the time it takes to move memories from one layer to another depends on the age of the memory, such that in early layers memories stay for a shorter amount of time than in later layers, this memory system could approximate a Weber-Fechner law, where the precision of determining the exact age of a memory decreases with its age. If something like a Age-Organization model is indeed implemented in some species, a fixed number of layers and the indirect pathways (gray arrows in Figure 6A) are probably established during development. An exact one-hot code for time, and an exact copy mechanism from one layer to the next seem unrealistic, but approximations thereof with bump-like time coding and linear transformations of the content from one layer to the next would lead to approximately the same behavior as the Age-Organization model.

#### 2.4.4 Age Organization with Synaptic Pruning

Another instantiation of a chronological organization model arises when considering the product between content and population-rate encoded “when” information (Figure 6B). This model is similar to the Poprate-Age-Tagging model. In both models, many synapses are grown during storage (storage at *t*_1_ in Figure 6B) and pruned at different moments in time, such that the age of a recalled memory can be decoded from the identities or numbers of active neurons during recall. The chronological organization across the memory system arises from the fact that synapses onto neurons in the first layer of the memory decay more quickly than those in the last layer (Figure 6B).

#### 2.4.5 Sparse Encoding with Random Synaptic Pruning and Simple Readout

Despite the frequent appearance of the special association schemes “concatenation” and “product” in the literature (Figure 2C), it is unclear why brains should favor them over other non-linear mixed association schemes.

Combining sparse random projections for association (Figure 2C) with synaptic delays for storage and recall (Figure 3) and synaptic pruning (Figure 5E and Figure 6B) leads to the Random-Pruning model (Figure 7). During storage, the input triggers distributed activity patterns in the intermediate and content neurons. Because of the random projections, these activity patterns encode the input information implicitly, i.e. the neurons in these groups are not necessarily “tuned” to a single feature, like the redness of an object, but a given neuron may specialize to specific combinations of features and become active, for example, only when a red triangle is shown. During the storage phase, a Hebbian plasticity rule can initiate the growth of synaptic connections between the intermediate layer and the content neurons (storage in Figure 7).

**Fig 7.**
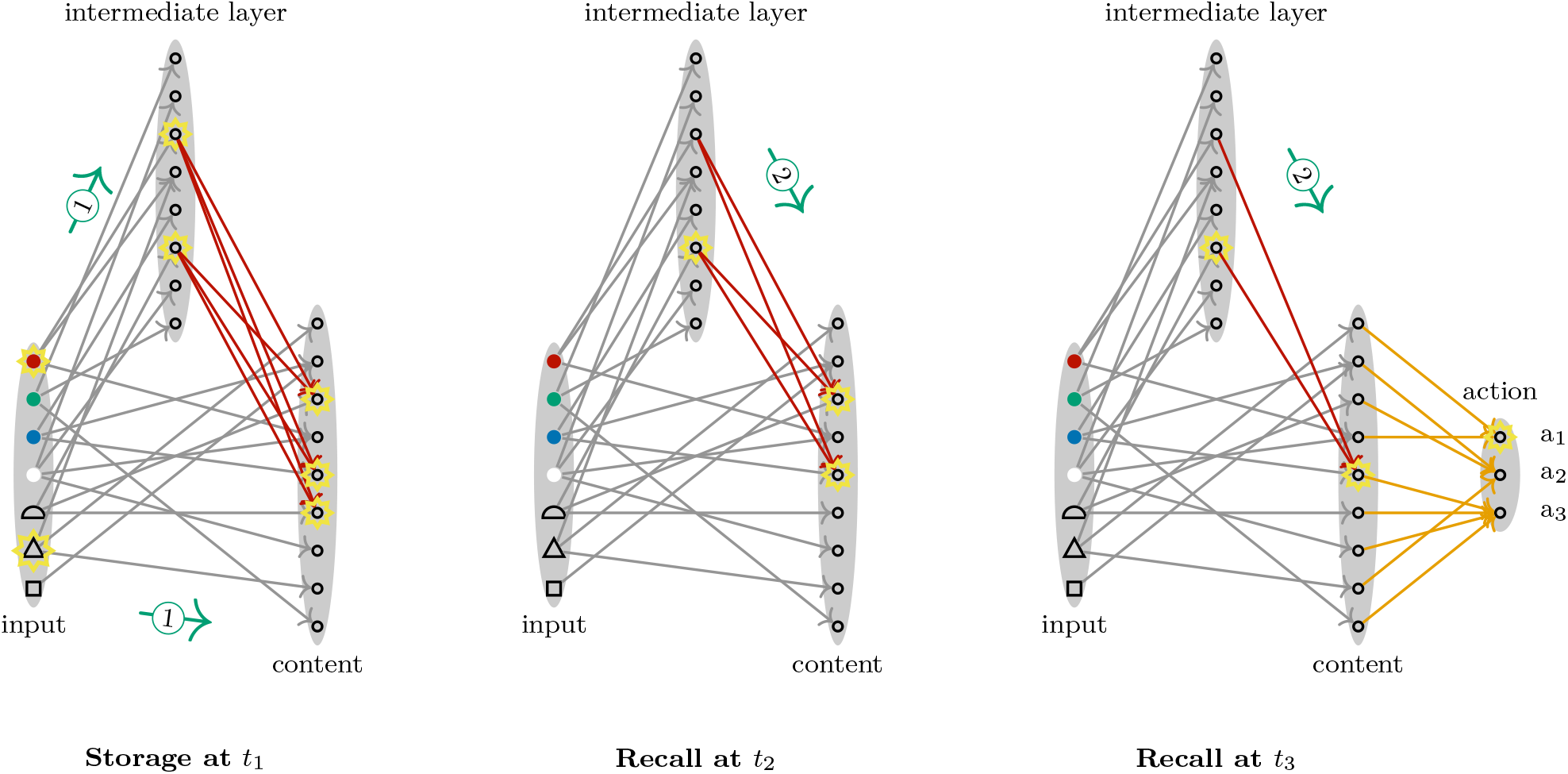
Sparse Encoding with Random Synaptic Pruning and Simple Readout. The input is sparsely and randomly connected to an intermediate and a content layer (gray arrows). During storage, the content and intermediate layers are driven by the input layer (green arrows with label 1), and Hebbian plasticity connects co-activated neurons (red arrows). Once the activity during recall with the triangle cue has reached the intermediate layer, the neurons in this layer drive the content layer through the red connections (green arrows with label 2; c.f. Figure 3) These synaptic connections are pruned after random, postsynaptic-neuron-specific durations, such that during recall at time *t*_2_ more neurons are activated in the content layer than during recall at time *t*_3_ *> t*_2_. This is an example of a non-linear mixed code of content and time, because the activity of a given neuron in the content layer can mean, for example, “a red triangle was observed at most so-and-so long ago”. The neurons in the content layer link directly to action neurons (orange arrows).

**Fig 8.**
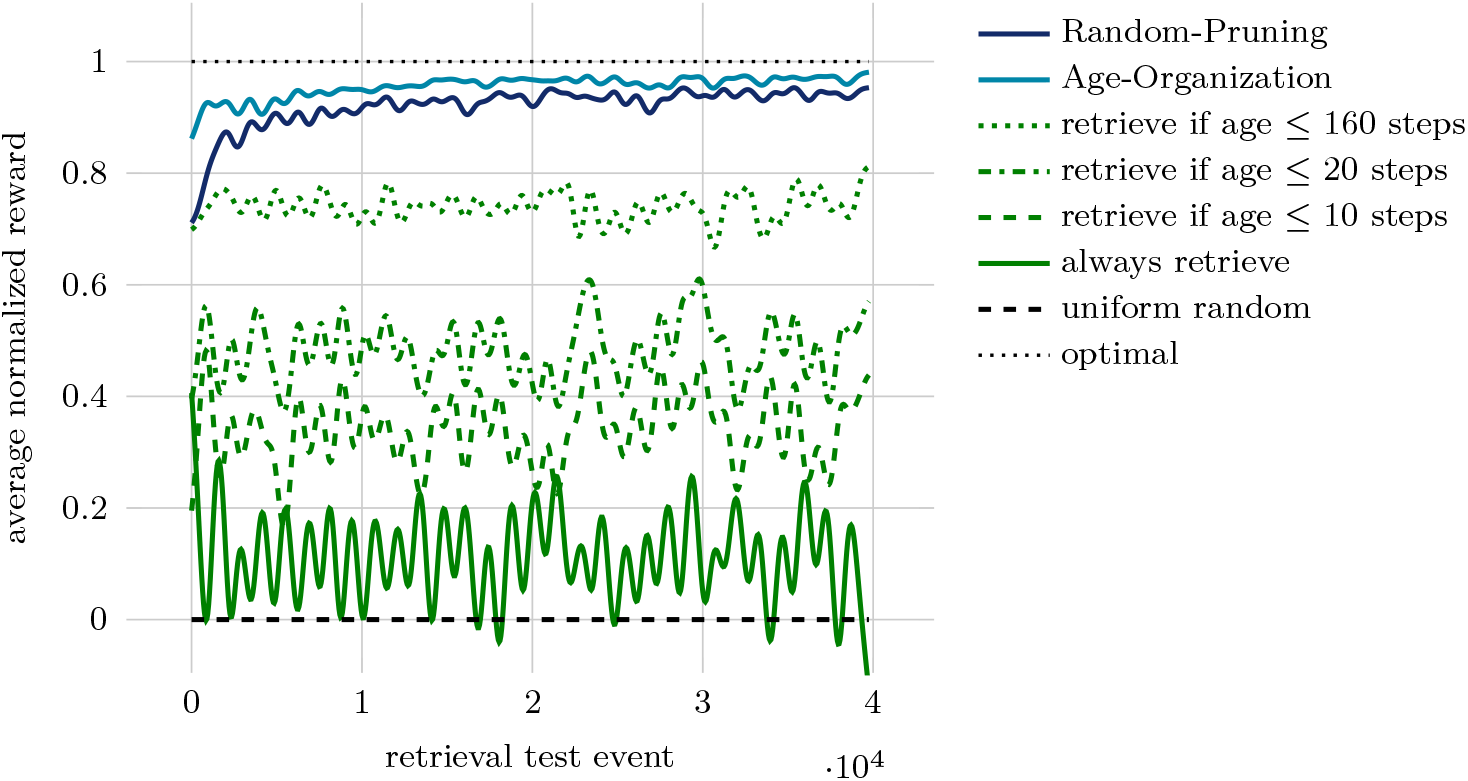
A scaled-up version of a variant of the Age-Organization and the Random-Pruning model yield good performance in a simulated food caching and retrieval setting. Average normalized rewards were computed with Gaussian smoothing (with standard deviation *σ* = 500 steps), and dynamic rescaling, such that the optimal policy always has reward 1, and uniformly random choices of actions *a*_1_ and *a*_2_ have reward 0. The performances of baseline policies (in green) are computed with full access to the history of caching events. Choosing always the retrieval action *a*_1_ (solid green line) incurs high costs for all locations, where the food item is not retrievable, whereas choosing the retrieval action only, if there was a cache event at the test location within the last 10 steps (dashed green line) misses many opportunities with retrievable items. The best baseline policy without food-type-specificity is to choose retrieval action *a*_1_ whenever at the given test location a food item was cached within the last 160 steps (dotted green line). The Age-Organization and the Random-Pruning model learn quickly to outperform the baseline policies and approach the optimal performance.

In the model of Figure 7, the sparse random projection code of the content is combined with a population rate code of the “when” information, similarly to the Poprate-Age-Tagging model (Figure 5E) and the chronological organization model with synaptic pruning (Figure 6B): synapses to the content layer are pruned after random durations that depend on the identity of the post-synaptic neuron, such that more neurons become active when recalling a recent event (recall at *t*_2_ in Figure 7), than when recalling an old event (recall at *t*_3_ in Figure 7).

#### 2.4.6 A What-Where-When Memory System for Food Caching Animals

As an example of where such a memory system could be beneficial in natural settings, let us examine food caching behavior (Figure 8). Scatter hoarders, such as nutcrackers or jays, store small food items in hundreds of locations each season [79]. There is good evidence that retrieval of their own caches is memory-dependent: olfactory cues are unnecessary; visual cues matter; and random search at preferred locations is inconsistent with the observed behavior [79]. As different types of food degrade at different rates, it is beneficial for these animals to keep memories not only of what they cached where, but also, how long ago the caching events happened. In the laboratory, *California scrub-jays* were observed to remember what they cached how long ago, and adapt their behavior, if they learn that certain types of food degrade faster or slower than others [33, 80–82].

To demonstrate that the memory systems proposed in this article enable good performance in these kinds of settings, we ran simulations with approximately 20’000 caching events of 3 different types of food at 1000 possible cache locations (Figure 8; in the abstract setting of Figure 1A this corresponds to 1000 distinct colors and 3 distinct shapes). Location and food type are given as two one-hot coded vectors to the input of the models. Randomly interleaved with the caching events, approximately 40’000 retrieval test events probed the memory system with possible cache locations. Response action *a*_1_ is interpreted as an active retrieval event at the test location (in the wild, a jay may start digging with its bill), whereas action *a*_2_ is interpreted as ignoring the test location (reducing waste of energy and time of an active retrieval event at a location where no food is cached or the food has already degraded). A positive reward for successful retrieval is given, if the retrieval action *a*_1_ is chosen at a location where food of the first type was cached less than 10 steps ago, or food of the second type less than 20 steps ago, or food of the third type less than 160 steps ago; a small negative reward for the wasted effort is given, if the retrieval action *a*_1_ is chosen otherwise; no reward is given for ignoring retrieval with action *a*_2_.

The Age-Organization model in Figure 8 relies on synaptic pruning after different delays as in Figure 6B. These delays are chosen artificially at 10, 20, and 160 steps, matching the degradation duration of the three types of food. At the beginning of the simulation, the readout weights are initialized such that action *a*_1_ is only chosen, when a caching event at the given test location happened less than 160 steps ago (best baseline condition in Figure 8), i.e. the network reacts to all food types the same. Within only a few tens of retrieval test events, a food-type-specialization is learned, and the memory system approaches the optimal policy. Conceptually related, but less artificial is the Random-Pruning model in Figure 8, which relies on sparse fixed random connections to an intermediate layer of almost 5000 neurons, sparse fixed random connections to a content layer of 60 neurons, and synaptic pruning after random, postsynaptic-neuron-specific durations in the range of 1 to 200 steps, as in Figure 7. The readout weights of this model are initialized such that action *a*_1_ is chosen, whenever a content neuron becomes active during recall, and action *a*_2_ otherwise. Although performing slightly worse than the artificial Age-Organization model specifically designed for this task, the Random-Pruning model performs considerably better than all baselines, and reaches close to optimal performance.

### 2.5 Experimental Predictions

To highlight advantages and disadvantages of different systems and explore the limitations of purely behavioral experiments as a tool to learn about how brains allow to remember the “when” of past events, we simulated different “what-where-when” memory models. For these simulations we assume that the subjects have to learn the behavioral rule through reinforcement learning, as is typically the case in experiments with animals.

For the Context-Tagging model we assume a fixed preprocessing to a one-hot intermediate representation of the age of a memory (Figure 4C), which is identical to the one-hot representation of time in the Onehot-Age-Tagging model. Because of their similarities, we do not simulate these two models separately and refer to them as Context/Onehot-Tagging model.

Although the discrimination of some models requires recordings of neural or synaptic dynamics, purely behavioral experiments can provide valuable insights. Suppose, for example, that a subject has learned how to respond to recalling a memory with a certain content and age, like performing action a_2_ when the event “red triangle” is remembered to have happened Δ*t*_train_ ago (Figure 9A, see also section “Protocols of Simulated Experiments”). In a similar task jays learned to avoid food caches containing crickets that they cached 4 days ago [24]. If one tests the subject on untrained content-age combinations, for example “blue square” after Δ*t*_test_ (Figure 9A), different representations and associations of content and memory make different predictions.

**Fig 9.**
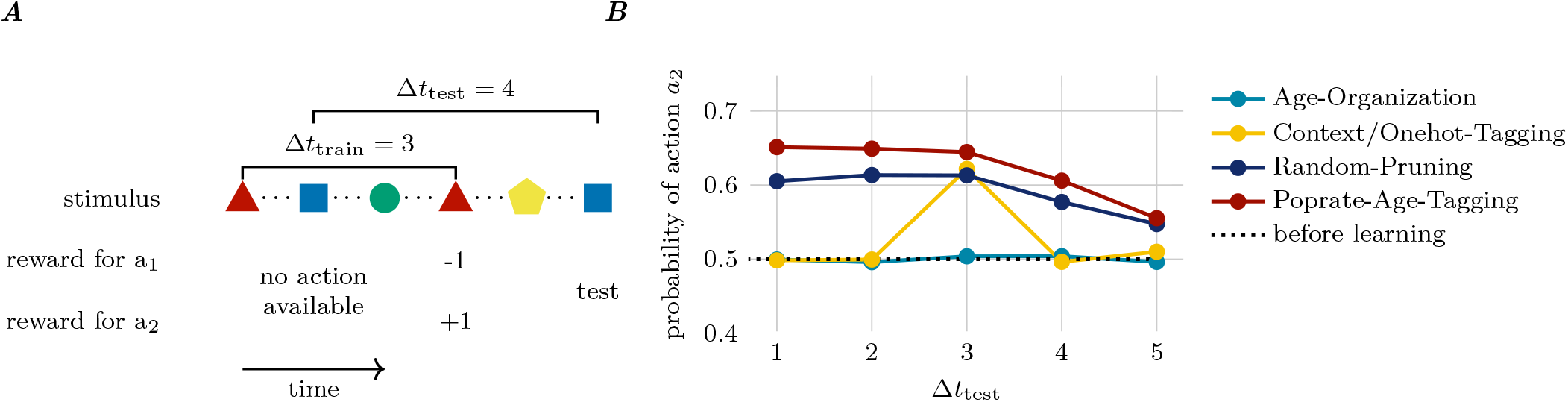
Generalization of actions that depend on the age of memories, based on a single rewarded trial. **A** A subject learns in a binary forced-choice task that action *a*_2_ is rewarded (or action *a*_1_ is punished), when recalling an event that happened Δ*t*_train_ ago. Training occurs with the same stimulus (red triangle). After training, the subject is tested once with a different stimulus (e.g. blue square) and a retention interval Δ*t*_test_ which may differ from the training interval. **B** Average probability across 10^4^ simulated subjects of taking action *a*_2_ as a function of Δ*t*_test_. For the sparsest code (Age-Organization) there may not be any generalization to other stimuli, even when the test interval is the same as the training interval (blue dot at Δ_test_ = 3). Conversely, for distributed representations (Poprate-Age-Tagging and Random-Pruning) there is generalization to other stimuli and test intervals different than the training interval. Quantitatively the results would be different for other learning rates or other values of the probability of *a*_2_ before learning, but qualitatively the results stay the same.

How a model generalizes depends mostly on the overlap of the recalled memories. For the one-hot coded memories in the Age-Organization model, there is no generalization from training to test settings, because distinct neurons are active during the recall of “red triangle Δ*t*_train_ ago” and “blue square Δ*t*_test_ ago” for any Δ*t*_test_ (light blue curve in Figure 9B). In the Context/Onehot-Tagging model, the tags for “red triangle Δ*t*_train_ ago” and “blue triangle Δ*t*_test_ = Δ*t*_train_ ago” are identical, despite the contents being different, and therefore there is some generalization to other memories of the same age (yellow curve in Figure 9B). Even more generalization occurs with the Poprate-Age-Tagging and the Random-Pruning model, because there is also some overlap in the recalled activity patterns for Δ*t*_test_ ≠ Δ*t*_train_.

Experiments that probe the learnability of different tasks can provide further evidence in favor or against specific models. Consider a task, where subjects are repeatedly trained to respond with action a_1_ if the age of a remembered event is less than some threshold and respond with action a_2_ otherwise (Figure 10A, see also section “Protocols of Simulated Experiments”). Such a task can be learned with all the models considered here, but the Age-Organization model has potentially an advantage, because the one-hot encoding permits faster learning with higher learning rates than other representations (Figure 10B; learning rates for all models are optimized for best final performance in the tasks in Figure 10A and Figure 11A). However, if an experiment would show slow learning, this should not be taken as evidence against the Age-Organization model, because a suboptimal learning rate would induce slow learning also in the Age-Organization model.

**Fig 10.**
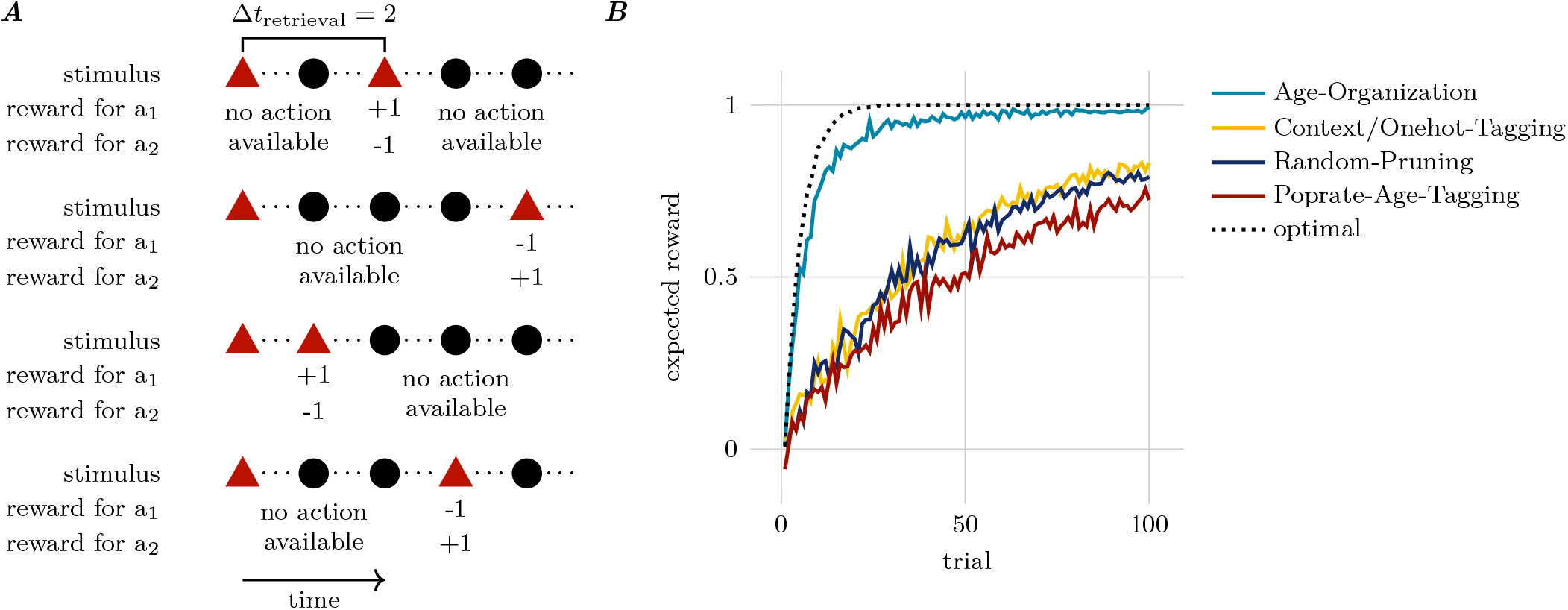
Learning to take actions that depend only on the age of memories. **A** The experiment consists of multiple trials with random retrieval intervals Δ_retrieval_. If the retrieval interval satisfies Δ_retrieval_ ≤ 2, action a_1_ is rewarded (+1) and action a_2_ is punished (reward −1); reward contingencies are reversed, if the retrieval interval is larger than 2. **B** The expected reward per trial is measured over 10^3^ simulated agents. All models can learn this task, but learning with one-hot codes can be faster than with other codes, because large learning rates can be chosen. The optimal performance (dashed line) was computed by averaging 10^4^ agents that make for each interval at most one mistake and always select the correct action afterwards.

**Fig 11.**
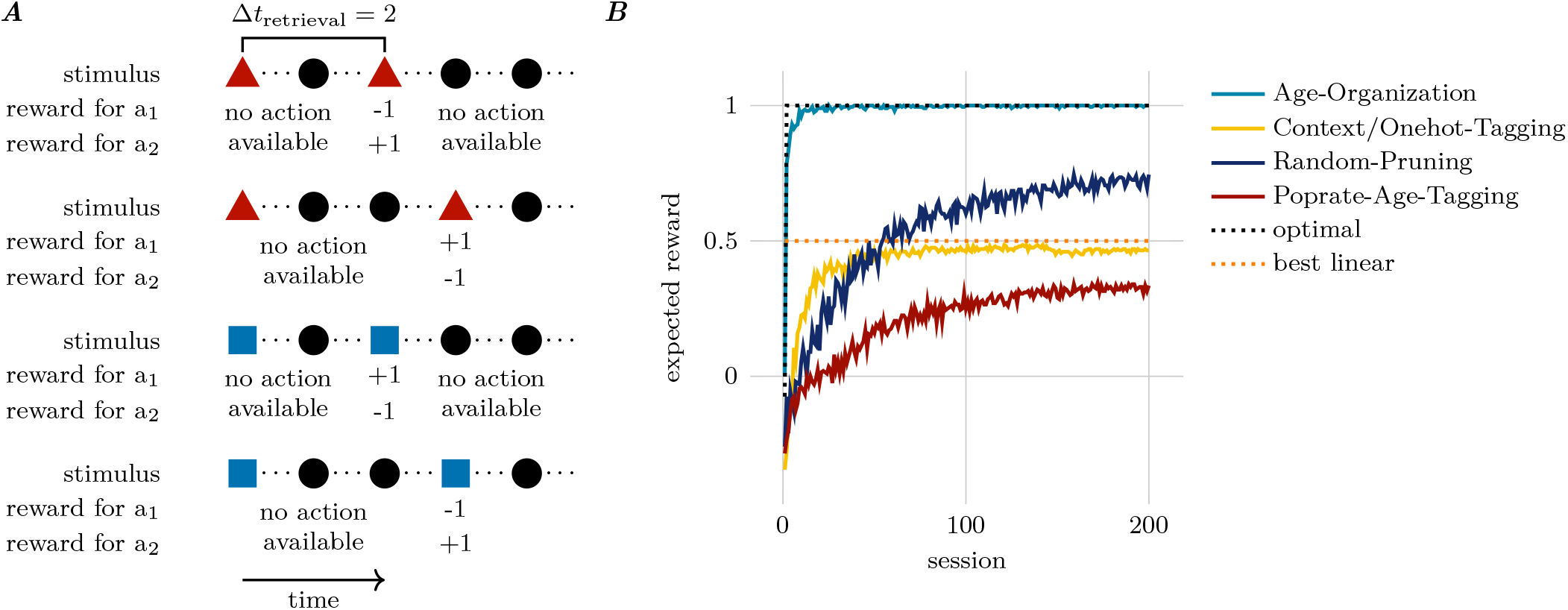
Learning to take actions that depend on the age and the content of memories. **A** The experiment consists of multiple trials with different retrieval intervals Δ_retrieval_ and objects. If the object is red and the retrieval interval is Δ_retrieval_ = 2 or the object is blue and Δ_retrieval_ = 3, action a_2_ is rewarded (+1) and action a_1_ is punished (reward −1); reward contingencies are reversed, otherwise. We call the sequence of these four trials one session. **B** The expected reward per session is measured over 10^3^ simulated agents. Because this is an XOR-like task, models with Context/Onehot-Tagging or Poprate-Age-Tagging encoding cannot reach better performance than the best linear model (correct in 3 and wrong in one condition leads to an expected reward of (+3 − 1)*/*4 = 0.5). The Random-Pruning model eventually learns the task, but it learns slower than the Age-Organization model encoding and sufficiently large learning rate. The optimal performance (dashed line) was computed by averaging 10^2^ agents that make for each interval and content at most one mistake and always select the correct action afterwards.

If the correct responses of the subjects depend not only on the age of the recalled events, but also on their content, XOR-like tasks can be constructed (Figure 11A, see also section “Protocols of Simulated Experiments”). For example, consider a task where action a_2_ is rewarded, whenever the memory of a red triangle is at most 2 time units old or the memory of a blue square is older than 2 time units, but otherwise action a_1_ is rewarded. A linear readout cannot correctly learn this rule, if the content and the “when” information are given in a concatenation code (Figure 11B). We find that tagging models systematically fail on this task (yellow and red curve in Figure 11B). Both the Age-Organization and the Random-Pruning can learn this task, but the Age-Organization model could learn it much faster (optimized learning rates, same as in Figure 10B).

The results in these simulated experiments depend crucially on the representation of the recalled information that arrives as input to the final linear decision making layer and the plasticity rule that changes the synaptic weights of this decision making layer. With this insight, it is straightforward to design similar experiments that investigate, for example, the representation of different aspects of “what” and “where” information. However, this dependence upon the representation of recalled information also implies that purely behavioral experiments cannot discriminate between different models that exhibit the same activity pattern as input to the final decision making layer. Therefore it may, for example, be almost impossible to discriminate timestamp from age representations of time, unless one can selectively manipulate the zeitgeber or rely on neural recordings.

## 3 Discussion

We showed that different choices of neural encoding, reference point of time, content-time associations, retrieval and readout mechanisms lead to a family of models, where the “what”, “where” and “when” of events can be stored and retrieved through automatic processes and Hebbian plasticity. Although the concrete implementations are idealized “toy”-models, that illustrate the central ideas succinctly, the same ideas generalize to bigger networks and more difficult tasks, as demonstrated in our simulation of food caching behavior.

The considered neural codes (rate, one-hot, distributed and poprate, Figure 2A) are spatial codes, in the sense that all relevant information about the “what”, “where” and “when” of an event is given by the activity pattern of a group of neurons in a single time step. This activity pattern could be, for example, the average firing rates of neurons in a time window of 100 milliseconds. In addition, one could consider spatio-temporal codes, where some information is encoded in the temporal evolution of activity patterns. For example, a single neuron could implement a temporal one-hot code, where the information is encoded by the duration between some fixed reference point in time and a spike (time-to-spike code). For spatio-temporal codes, more sophisticated readout and learning mechanisms than the ones in Figure 2F would be needed to extract information from the temporal evolution of activity patterns. With spatio-temporal codes, the already large lower bound of 288 models (see section “Experimental Predictions”) would further increase and include models with spatio-temporal retrieval (e.g. [83]).

Chronological organization models can be generalized to include models where the different groups of content neurons are not just copies of one another, but the representations of the memory content in each group differ from on another. For example, a lossy, age organized model, similar to the one described in section “Age Organization with Synaptic Pruning”, could store the gist of an event in some groups of neurons together with a detailed representation in other groups of neurons and forget the detailed representation faster than the gist. This could be a simple model of the trace transformation theory [16], which postulates that recall of details requires a functional hippocampus, whereas the gist can be recalled without hippocampus.

On a conceptual level, multiple theories of the processing of “when” information have been proposed. For example, [10] described eight theories: strength, chronological organization, time tagging, contextual overlap, encoding perturbation, associative chaining, reconstruction and order codes. The last four theories of this list are beyond the scope of this article. Our work, however, provides concrete hypotheses for neural implementations of the first four theories and discusses their implications on readout of temporal and content information.

Although we focused on long-term memory with behavioral readout, we briefly summarize here known results where storage and retrieval are separated by less than a few minutes, including those with human subjects where the behavior is instructed. In psychology, there is a huge literature on laboratory-based memory tasks on short timescales [84]. Among the most popular computational models to explain these memory experiments are different variants of temporal context models [18, 64–66] (see also Table 2), which keep memories of the recent past with decaying activity traces (temporal context) and learn associations between temporal context and individual memory items with fast synaptic changes. These models can thus be seen as implementing both a rate code of age information and a distributed timestamp code (Table 2). Whereas the decaying activity traces are unlikely to extend to timescales of days or years, there is some experimental support for the contextual overlap theory also for long timescales: for example, recency judgments were found to be context-dependent [85]; furthermore, the contiguity effect of remembering multiple events jointly, if they happened at similar moments in time, seems to generalize to autobiographical memory and can be explained with temporal context models [86]. However, there is also evidence for hippocampus-dependent reconstruction-based theories in humans [87].

The richness of biological phenomena in general, and memory phenomena in particular, in combination with the “theory-ladenness” of observations [88], allows for finding support in experimental data for different theories. Memory storage in the proposed models relies on one-shot Hebbian synaptic plasticity, for which there is experimental evidence [57, 89]. Simple reinforcement learning of new tasks requires neoHebbian synaptic plasticity (see section “Synaptic Plasticity”), where plasticity depends on a modulatory signal that arrives up to a few seconds after Hebbian tagging; there is ample experimental evidence for this kind of plasticity [45–49]. Age representations of time in synaptic memories requires ongoing changes of synaptic strengths or connections. An implementation of age representations of time with a rate code could rely on synapses that decay at different rates on a timescale of days to weeks [90, 91]. One-hot or distributed encoding of the age of memories requires continual growing and pruning synapses. Synaptic turnover has been widely observed [58–62, 92], but we are not aware of experimental results that would support or refute the rewiring mechanisms proposed in our models. The specific pruning mechanism involving neuron-dependent pruning rates on a timescale from hours to years (Synaptic Plasticity) is not inconsistent with current knowledge of synaptic turnover, but direct experimental evidence for this hypothesis is lacking. Ongoing synaptic changes are also required in other theories and models of systems memory consolidation [16, 93], and could potentially be mediated with mechanisms such as parallel synaptic pathways [76] or memory transfer [75]. Although there is experimental evidence – of varying strenght – for all the synaptic processes needed to implement the above models, further experiments are clearly necessary to determine which processes are actually used by different species to remember the time of past events.

On timescales from a few hundred milliseconds to tens of seconds, and potentially up to a few hours, activity-based age tracking of memories may be mediated by time cells [9, 69–74]. Time cells are neurons that fire at successive moments in temporally structured experiences. They were found in region CA1 of the hippocampus of rodents and in the entorhinal cortex of macaque monkeys [94] (but see [95], for an example where no time cells were found in CA1). Time cells could be seen as evidence for activity-based models of age tracking with concatenation or product associations (see TCM or TILT model in Table 2), but a random spatio-temporal feature model, akin to the Random-Pruning model, would probably lead to similar results. XOR-like experiments, like the one suggested in Figure 11, could potentially be used to distinguish these hypotheses for age tracking on small timescales.

More relevant for the topic of this paper are experiments involving longer timescales. For example, neural responses to the same context in the CA1 region of the rodent hippocampus have been shown to drift over hours and days [96–100]. If the recorded activity in these studies reflects a temporal context tag used for memory storage, such slow representational drift would be consistent with a timestamp mechanism similar to the Context-Tagging model. On the other hand, if the measured activity primarily reflects the recall of a salient past experience – such as the first exposure to that context – these findings would favor an age-coding interpretation of past events. While it appears less likely that recall dominates the recorded activity, further experiments that clearly separate storage and recall phases are necessary to distinguish between these interpretations. Lastly, it is also possible that the observed neural activity is unrelated to the storage or recall of specific events and instead reflects factors such as behavioral variability [101].

Sparse codes enable fast and flexible learning of behavioral rules, as demonstrated by the Age-Organization model. While explicit one-hot coding of content and time with single neurons does not seem plausible, relaxed versions of our models with redundant, sparse, and almost not overlapping codes are consistent with experimental findings of sparse activity patterns in granule cells of the dentate gyrus (DG) of rodents [102–104] or the recently discovered “barcode” activity patterns in the hippocampus of food-caching birds [105]. Sparse activity patterns in the intermediate layer of the Random-Pruning model (Figure 7) were also crucial to scale-up our simulations (Figure 8). Models with sparse codes get also indirect support from behavioral studies: experiments with *California scrub-jays* showed convincingly that these birds have a flexible “what-where-when” memory system [14, 22–25, 106]. Although the existing experimental results cannot discriminate between the different models considered here, the observation that they can learn different behavioral rules that depend on the content and age of memories within few trials, speaks in favor of an Age-Organization model or a Random-Pruning model, that allow fast and flexible learning. To get more behavioral evidence for the different models, the suggested experiments in Figure 9, Figure 10 and Figure 11 could be run in the similar spirit as these food caching experiments or the non-verbal experiments with other species mentioned in Table 1.

Remembering the “when” is an idiosyncratic feature of episodic and episodic-like memory. Thus, revealing the mechanisms that underlie the ability to estimate the age of memories is a crucial step towards a better understanding of episodic memory systems. The different models discussed here can serve as concrete hypotheses and the simulated experiments as inspirations for future experiments that combine behavioral and physiological recordings to learn more about how humans and animals remember the “when” of past events.

## 4 Methods

### 4.1 Formal Description of Codes

We consider a discretized version of the sensory stream in 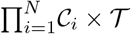, where 𝒞_*i*_ denotes the finite set of values that sensor *i* can take and 𝒯 denotes the finite set of possible time points.

For a single finite set 𝒞= {*c*_1_, …, *c* _|𝒞 |_}with elements *c*_*i*_ and cardinality |𝒞 |, we define three different elementary coding schemes for representing element *c*_*i*_:

- *rate code*: `rate`(*c*_*i*_) := *r*(*i*) ∈ ℝ, where *r* is an arbitrary, one-to-one function.
- *one-hot code*: a |𝒞 |-tuple-valued function `onehot`(*c*_*i*_) := `onehot`_1_(*c*_*i*_), …, `onehot`_|𝒞 |_ (*c*_*i*_) with `onehot`_*j*_(*c*_*i*_) := *A*_*i*_*δ*_*ij*_, amplitude *A*_*i*_ *>* 0 and Kronecker delta *δ*_*ij*_ := 1 if *i* = *j* and *δ*_*ij*_ := 0 otherwise.
- *distributed code*: a one-to-one *M*-tuple-valued function or random vector `distr`(*c*_*i*_) := `distr`_1_(*c*_*i*_), …, `distr`_*M*_ (*c*_*i*_) with *M >* 1 and 0 *<* `distr`_*j*_(*c*_*i*_) ≤ `distr`_*j*′_ (*c*_*i*_) for at least one pair *j* ≠ *j*′ and at least one *i* (Figure 2A).

In addition to these elementary coding schemes, we consider population rate codes, where the value *c*_*i*_ is encoded by the number of active neurons. An example of a deterministic population rate code is given by `distr`_*j*_(*c*_*i*_) = 1 if *j* ≤ *i* and `distr`_*j*_(*c*_*i*_) = 0 otherwise (`poprate` in Figure 2A). Stochastic population rate codes satisfy the condition Pr (∑_*j*_ `distr`_*j*_(*c*_*i*_) = *i*) = 1. These codes can be reduced to a rate code by computing the population activity rate 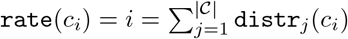.

We write *ℓ*(`code`(𝒞)) or *ℓ*(`code`) for the length of an element of in `code`-coding, e.g. *ℓ*(`rate`(𝒞)) = 1, *ℓ*(`onehot`(𝒞)) = . |𝒞 |

A generalization to continuous variables *x* (space, time, color, etc.) can easily be found with, e.g. continuous rate coding `rate`(*x*) = *x*, generalized one-hot coding with e.g. radial basis functions 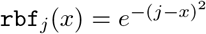 for a population of neurons with indices *j* = 1, …, *M* or generalized distributed code with e.g. mixtures of radial basis functions.

### 4.2 Formal Description of Association Schemes

Any function 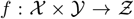 defines an association 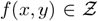 between elements *x* ∈ 𝒳 and *y* ∈ 𝒴. If the function *f* is one-to-one, the association *f* (*x, y*) keeps the full information about the associated elements *x* and *y*.

We define

- *concatenation of codes*: ***x*** ⊕ ***y*** := (*x*_1_, …, *x*_ℓ (*x*)_, *y*_1_, …, *y*_ℓ (*y*)_) (Figure 2C).
- *linear mixed codes*: ***x*** ⊕_*M*_ ***y*** := *M* (***x*** ⊕ ***y***), where *M* : ℝ^ℓ(***x***)+ℓ(***y***)^ → ℝ^*M*^ is a linear map.
- *product of codes*: ***x*** ⊗ ***y*** := ***x*** · *y*_1_ ⊕ ***x*** · *y*_2_ ⊕ · · · ⊕ ***x*** · *y*_ℓ(*x*)_, where the product ***x*** · *y*_*i*_ between a tuple ***x*** and a scalar *y*_*i*_ has the standard meaning ***x*** · *y*_*i*_ = (*x*_1_*y*_*i*_, …, *x*_ℓ(*x*)_*y*_*i*_) (Figure 2C)
- *random projections*: RP(***x, y***) = *σ*(*W*_1_***x*** + *W*_2_***y***), where *W*_1_ and *W*_2_ are fixed random matrices and *σ* is some non-linear function that is applied element-wise.

### 4.3 Mathematical Description of the Models

We model brains that observe sensory states ***x***_*t*_, ***x***_*t*+1_, … and take actions (or decisions) *a*_*t*_, *a*_*t*+1_, … on a slow timescale. These brains have internal neural states ***z***_*τ*_ and synaptic connection parameters *W*_*τ*_ that evolve on a faster timescale, indicated by the time index *τ* .

For all the models we used the storage and recall mechanism with synaptic delays, described in Figure 3 and hetero-associative recall. For the tagging models, this implementation differs from the descriptions with auto-associative recall in Figure 4 and Figure 5, but it leads to the same predictions for the behavioral experiments.

The internal neural states ***z***_*τ*_ are organized into groups of neurons. Tagging models have sensor, intermediate, content, tag and actuator neurons and we write the neural state 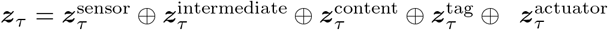. The Age-Organization model and the Random-Pruning model have the same groups of neurons, except that the group of tag neurons is lacking and the group of content neurons is larger than in the tagging models. The group of sensory neurons is further divided into two subgroups that receive color and shape as one-hot coded input. The activity in these sensory neurons propagates along the synaptic connections to down-stream neurons, until an action is taken. Once an action was taken, the next sensory input is provided to the sensory neurons. The activity propagation along synaptic connections can be described in terms of the update of the neural state of neurons in group *µ*, which is given by

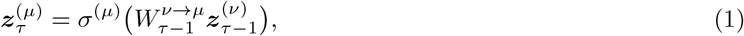

where *σ*^(*µ*)^, the activation function of neurons in group *µ*, is applied element-wise to the matrix-vector product of synaptic weight matrix 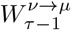 and activity state 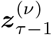 of group *ν* in the previous time-step *τ* − 1. The activation function of the actuator group is the soft-max function 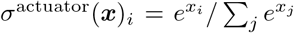. For all other groups of neurons we use the Heaviside function *σ*^(*i*)^(***x***)_*i*_ = *H*(*x*_*i*_ − *b*) = 1 if *x*_*i*_ *> b* and *H*(*x*_*i*_ − *b*) = 0, otherwise, where bias *b* = 0 for all groups of neurons except for the content group in the Random-Pruning model, where *b* = 1.5. The non-zero bias in the Random-Pruning assures sparse activity in the content layer. Action *a*_*t*_ is sampled with probability 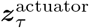 after recall has happened.

Because we used the recall mechanism with synaptic delays (Figure 3) and it takes three time steps for sensory activity to propagate along the “input-intermediate-content-actuator” pathway, there is a simple relationship between the slow timescale indexed by *t* and the fast timescale indexed by *τ* : if 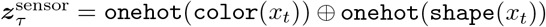, action *a*_*t*_ will be sampled with probability 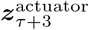. The sensory neurons are inactive during propagation of the activity through the neural network, i.e.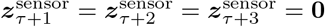; the sensory neurons are reactivated, once action *a*_*t*_ has been taken, i.e.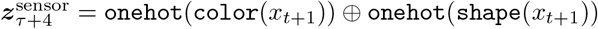.

Synaptic weight matrices are static or evolve according to one of the following plasticity rules:

#### Hebbian

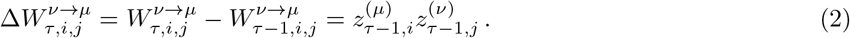

where *i* is the postsynaptic neuron in group *µ* and *j* the presynaptic neuron in group *ν*.

#### Reward-Modulated Hebbian

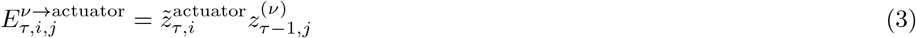

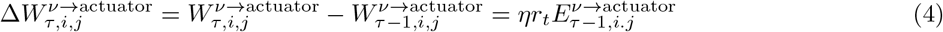

where 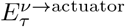 is an eligibility trace that depends on 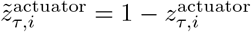 for postsynaptic neuron *i* = *a*_*t*_ and 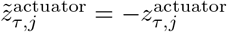, otherwise, *η* a learning rate and *r*_*t*_ ∈ {−1, 1} is the reward obtained after performing action *a*_*t* −1_. This plasticity rule can be seen as a policy gradient (REINFORCE) rule [107], where the terms 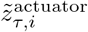 arise as a consequence of taking the derivative of the logarithm of the soft-max policy function in the derivation of the REINFORCE rule. This plasticity rule is used in all models for the readout weights (orange in the figures). We did not extensively optimize the learning rates, but ran the experiments in Figure 10 and Figure 11 for *η* ∈ {0.005, 0.01, 0.02, 0.05, 0.1, 0.2, 0.5, 1.0, 2.0 }, and picked for each model a value that lead to good performances on both tasks (Age-Organization: *η* = 1.0; Context/Onehot-Tagging: *η* = 0.5; Random-Pruning: *η* = 0.02; Poprate-Age-Tagging: *η* = 0.2). We used the same learning rates in all simulations reported in Figure 8, Figure 9, Figure 10, Figure 11.

#### Hebbian Latent-State-Decay

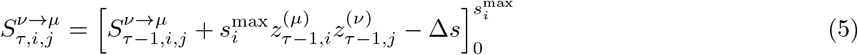

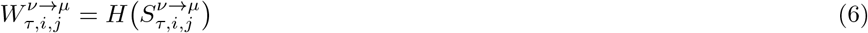

where 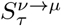 is a latent synaptic state matrix, 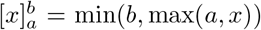, 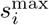 is the maximal value the latent state for post-synaptic neuron *i* can achieve, Δ*s* is a decay term and *H* is the Heaviside function. After a Hebbian growth, synapses of this kind are pruned after 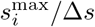 time-steps, unless they are restrengthened meanwhile. This plasticity rule is used in the Poprate-Age-Tagging model for the connections from intermediate to tag neurons and in the Random-Pruning model for connections from intermediate to content neurons. In the Poprate-Age-Tagging, we set 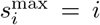 for the six tag neurons. In the Random-Pruning, we sampled 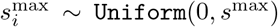, where *s*^max^ = 6 for the simulations in Figure 9, Figure 10 and Figure 11, and *s*^max^ = 200 for the larger model in Figure 8. We set Δ*s* = 1*/*3 for all models and simulations, implying that the latent states decay by one, for one slow time step indexed by *t*.

#### Postsynaptic Rewiring

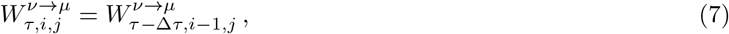

where Δ*τ* is such that the synaptic change happens always when perceiving a new input *x*_*t*_. This rule is an abstract implementation of a hypothetical systems consolidation process, where the active neuron encodes the age of a memory (e.g. Figure 4C or Figure 6A). This plasticity rule is used in the Onehot-Age-Tagging model for the connections between the intermediate neurons and the tag neurons and in the Age-Organization model for the connections between the intermediate neurons and the content neurons.

### 4.4 Equivalence of One-Hot coding and Deterministic Population-Rate coding

Let ***x*** be a one-hot coded neural activity pattern, ***w*** ∈ ℝ^*N*^ a weight vector, *y* = ***w***^*T*^ ***x*** a linear readout, and 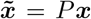, with *P* such that 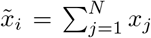, the re-parametrization from one-hot coding to the deterministic population rate code at the bottom of Figure 2A. The corresponding transformation 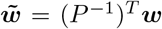 leaves the response invariant, i.e. 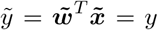. The gradient descent learning rule 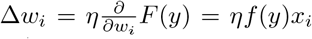, for some 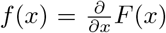, transforms under *P* to 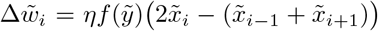.for *i* = 2, …, *N* − 1 and 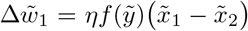, 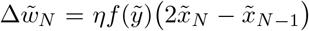 as can be seen by computing (*P*^−1^)^*T*^ (*P*) ^−1^ [108]. If the synapses are spatially organized such that the inputs of presynaptic neurons *i* − 1, *i, i* + 1 are neighbouring, the resulting plasticity rule features cross-talk between neighbouring synapses. In particular, a synaptic weight 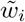 should only be changed, if the inputs at the neighbouring synapses *i* − 1 and *i* + 1 differ from the input at synapse *i*.

### 4.5 Protocols of Simulated Experiments

The experimental protocols of the simulated experiments are reported from the perspective of the experimenter. All protocols are constructed using four basic actions that an experimenter performs: show to the subject some stimulus (**`show_to!`**), provide a choice of multiple actions and observe the action taken by the subject (**`force_to_choose_and_observe_action!`**), reward or punish the subject (**`reward!`**) and keep track of the relevant observations in the results table (**`push!`**). It is assumed, but not explicitly modelled, that in between each action taken by the experimenter there is some waiting time. These waiting times should be sufficiently long, for example on the order of hours or days, such that subjects cannot solve the tasks with working memory alone, but need to rely on long-term “what-where-when” memory.

#### 4.5.1 Generalization of actions (Figure 9)

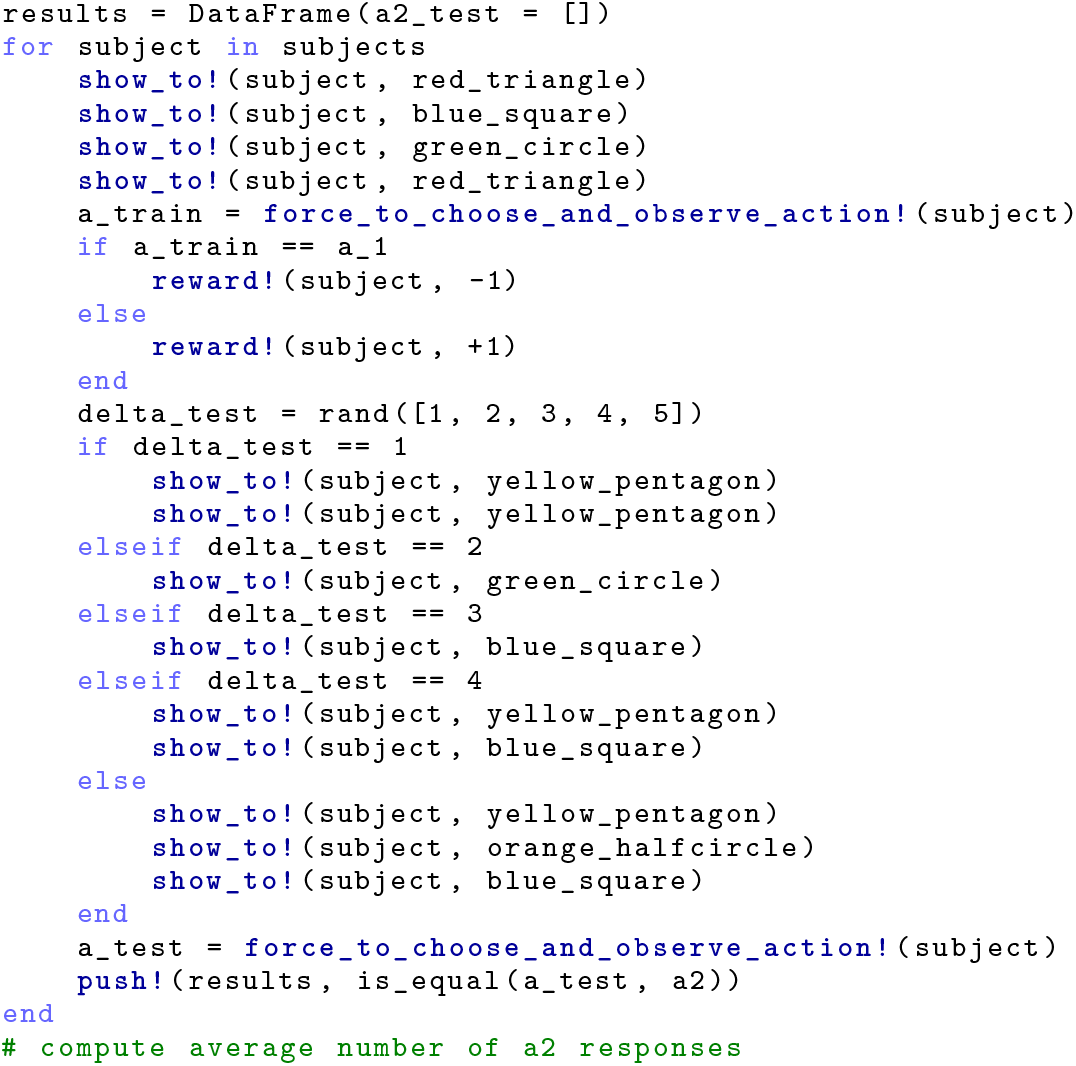

#### 4.5.2 Learning to take age-dependent actions (Figure 10)

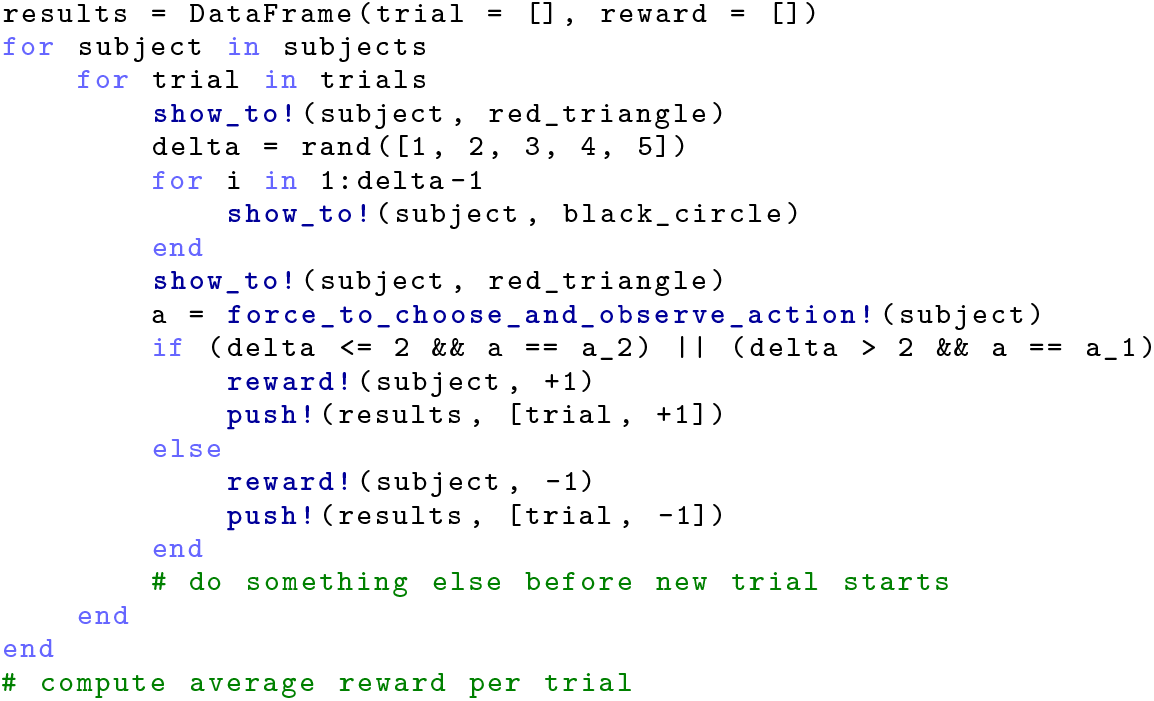

Learning to take age- and content-dependent actions (Figure 11)

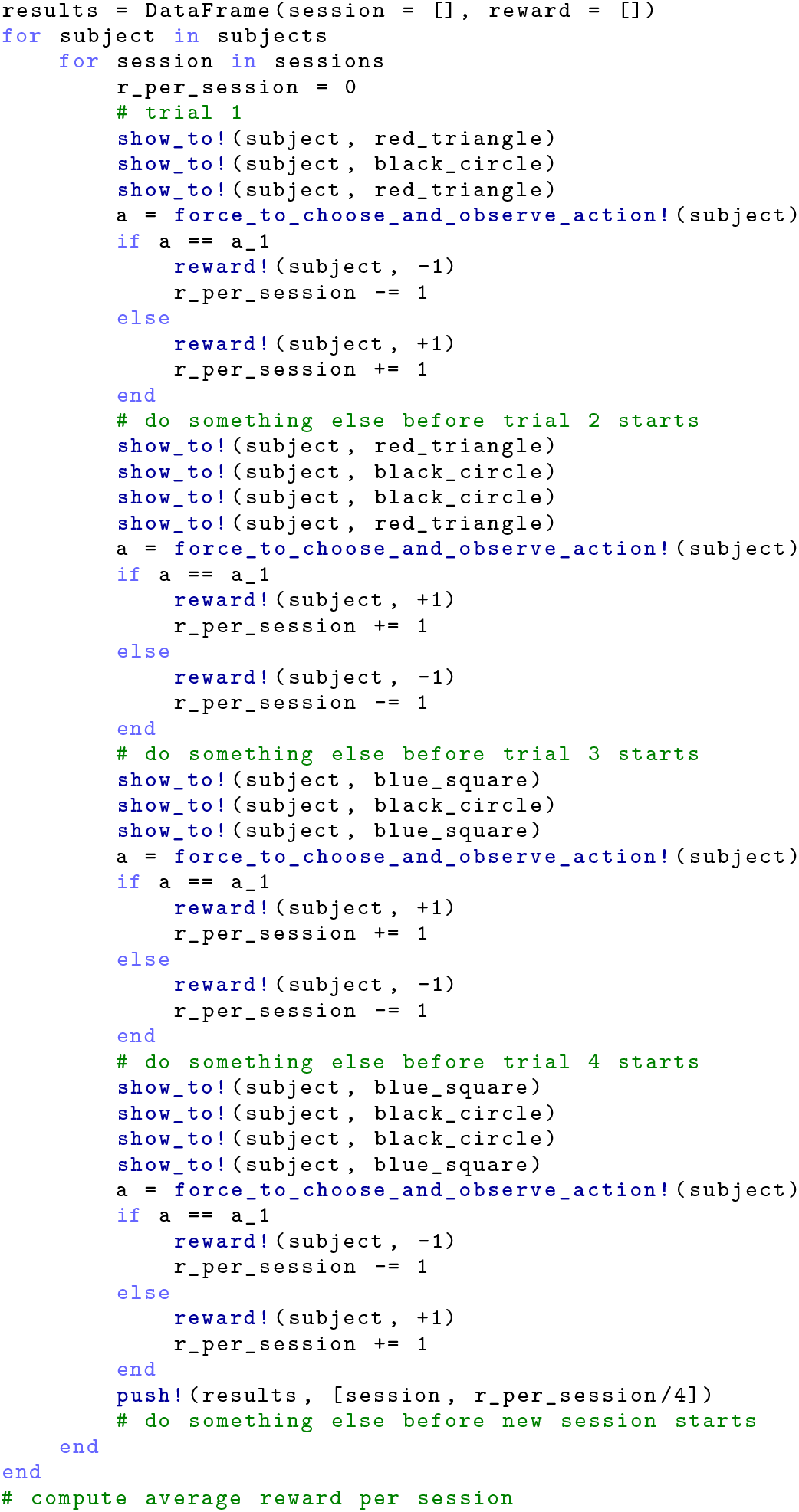

